# SPACE-Tag Enables High-Throughput Spatial Chromatin Profiling With CUT&Tag

**DOI:** 10.64898/2026.11.06.764689

**Authors:** Chao Yan, Ruiyang He, Sara Fernandez, Maroš Pleška, Silas Maniatis, Ivan Raimondi, Sanja Vickovic

## Abstract

Spatial epigenomics maps chromatin states within intact tissues, yet current methods remain limited by accessibility and scale. We developed SPACE-Tag, a spatial CUT&Tag framework that converts epigenetic marks-associated DNA into polyadenylated RNA through T7-based linear amplification, enabling direct integration with standard spatial transcriptomic workflows. Applied to the mouse brain, SPACE-Tag resolved activating and repressive chromatin landscapes, recovered region-specific gene-regulatory programs across anatomical domains, and showed strong correlation with existing bulk and single-cell reference data. SPACE-Tag enabled generation of an integrated multimodal atlas of the aging mouse brain, profiling five histone modifications and the transcriptome in consecutive tissue sections across the adult lifespan. This analysis uncovered sharp regional divergence with aging in the brain: fiber tracts showed widespread transcriptional downregulation, often without nearby chromatin state transitions, whereas the isocortex showed coordinated chromatin–transcriptome changes, including an H3K4me1 peak module that increased with age, was enriched at super-enhancers and linked to upregulation of neuronal plasticity regulators. These findings highlight the power of multimodal and high-throughput spatial genomics in uncovering the divergent regulatory logic that governs tissue-specific vulnerability and resilience in the aging brain.

---

By capturing transcriptional snapshots of cells *in situ*, spatial transcriptomics technologies have empowered exploration of cellular programs involved in tissue organization, intercellular communication, development, aging and disease^1–7^. In contrast to transcriptomic methods, the development and widespread adoption of methods for spatially resolved epigenetic profiling has been limited. Methods built on custom microfluidic barcoding have provided early proof of principle for spatial epigenomic profiling^8–12^, yet their reliance on specialized fabrication and relatively small capture areas make profiling intact tissues across multiple modifications and timepoints impractical, which hinders the spatiotemporal, multi-modal studies needed to resolve how chromatin landscapes evolve across organs and time. Other approaches, such as spatial ATAC-seq^13,14^, which were adapted for use with commercial platforms, can efficiently profile chromatin accessibility but do not directly resolve distinct histone modifications, thus providing only a partial view of the regulatory state. Finally, available approaches are often limited by sensitivity, specificity, and scalability, which restricts their ability to generate comprehensive maps across large tissues or study design cohorts.

To bridge the gap between cell-level chromatin assays and tissue-scale spatial profiling^15,16^, we developed SPACE-Tag, a spatial epigenomics method designed to address three key challenges in mapping histone modifications *in situ*: scalability, specificity and sensitivity (**Fig. 1a**). First, to enable scalable spatial measurements across large tissue sections, SPACE-Tag is implemented on the widely accessible Visium (10x Genomics) spatial array platform. Second, to achieve high molecular specificity for histone modifications, the method builds on CUT&Tag (Cleavage Under Targets and Tagmentation)^17^, in which antibody-guided protein A–Tn5 transposases selectively target defined chromatin marks. Third, to overcome the signal sparsity that often limits spatial epigenomic profiling, SPACE-Tag incorporates engineered adapters containing a T7 promoter that enables *in situ* linear amplification of genomic DNA fragments at sites of tagmentation ^18–21^. These adapters also include a poly(dT) tail that enables efficient capture on spatial transcriptomics arrays^15,16,22–24^, thereby linking amplified chromatin fragments into spatially indexed RNA molecules.

**Figure 1.**
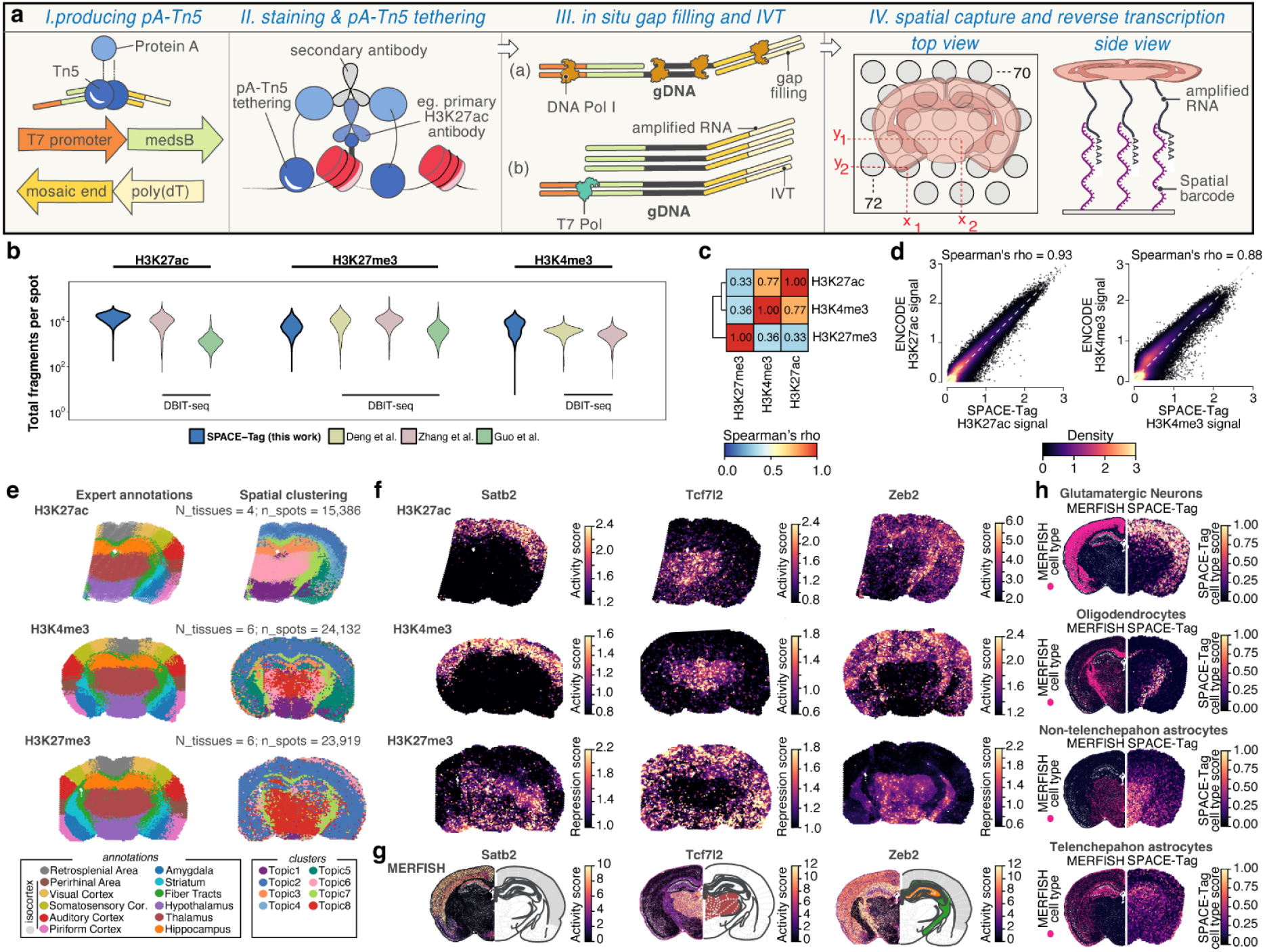
SPACE-Tag method overview and performance. (**a**) Schematic overview of the SPACE-Tag workflow. Fresh frozen tissue section is first mounted onto a solid-phase capture spatial transcriptomics slide. Following steps are then performed: I) Engineered transposome adaptors containing a T7 promoter and a poly(dT) sequence are loaded onto pA-Tn5. II) Primary antibody, secondary antibody, and pA-Tn5 are sequentially applied. III) Tagmentation and gap filling are carried out, followed by *in vitro* T7 transcription (IVT) to generate amplified RNA fragments. IV) The IVT-generated RNA is captured by poly(dT) probes, reverse transcribed, and processed into sequencing libraries. (**b**) Per-spot unique fragment counts (y-axis) obtained by SPACE-Tag and other published spatial chromatin profiling methods (x-axis). Spatial sampling method denoted for approaches based on DBIT-seq^10–12^. (**c**) Correlation matrix (color scale; Spearman’s rho) of histone marks profiled by SPACE-Tag. (**d**) Benchmarking SPACE-Tag against ENCODE. Scatter plots comparing SPACE-Tag (x-axis) and ENCODE (y-axis) reference datasets for H3K27ac (left) and H3K4me3 (right). Density (color scale) indicates the frequency of genomic peaks. The gray dashed line indicates the linear regression line. (**e**) Expert-annotated brain regions (left) compared with spatial maps derived from unsupervised topic clustering of histone modification profiles (right). Each row shows a different histone mark: H3K27ac (top), H3K4me3 (middle), and H3K27me3 (bottom). (**f**) Spatial projections of inferred gene activation (for H3K27ac and H3K4me3) or repression (for H3K27me3) (color scale). Columns show representative genes (left to right: *Satb2*, *Tcf7l2*, *Zeb2*), while rows show different histone modifications: H3K27ac (top), H3K4me3 (middle), and H3K27me3 (bottom). (**g**) MERFISH gene expression (color scale) for representative genes (left to right: *Satb2*, *Tcf7l2*, *Zeb2*). For each gene, the image pair shows MERFISH expression data from the Allen Brain Atlas (left) and the illustrated Allen Brain Atlas spatial regions where the gene is expected to be expressed (right). Spatial region color code is shared with expert annotations in (**e**). (**h**) Spatial cell type projections (color scale) in MERFISH (left) and SPACE-Tag (right) data. Four cell types (columns) are highlighted (from left to right: inhibitory neurons, oligodendrocytes, non-telencephalon astrocytes, and telencephalon astrocytes).

Consistent with this design, the T7 amplification step increased the local copy number of targeted fragments by approximately 55-fold while avoiding the amplification bias typical of exponential PCR^25^ (**Methods**, **Supplementary Fig. 1**). Additional optimization of tissue processing and enzymatic reactions, including fixation, nuclei permeabilization, antibody staining, Tn5 tagmentation and gap filling, further increased library complexity by approximately eight-fold (**Methods**, **Supplementary Fig. 2**). Together, these improvements provide sufficient sampling depth to resolve chromatin landscapes across diverse tissue domains. Using this approach, we examined region-specific reconfiguration of chromatin states during brain aging and investigated how spatial epigenomic patterns relate to transcriptional programs across grey and white matter.

To evaluate the sensitivity and specificity of SPACE-Tag, we profiled two activating (H3K27ac and H3K4me3) and one repressive (H3K27me3) histone modification in the mouse brain (N_animals = 6, N_sections = 16, and N_spots = 63,437). Sensitivity was assessed using the per-spot unique fragment count, while specificity was evaluated using the fraction of reads in peaks (FRiP), correlation across different modifications, and orthogonal benchmarking against chromatin accessibility. SPACE-Tag captured approximately ∼15,000 fragments per spot for H3K27ac, ∼6,000 for H3K4me3, and ∼6,000 for H3K27me3 (**Fig. 1b**), matching the performance of previously published spatial approaches^10–14^ while simultaneously yielding high FRiP values (0.78, 0.66, and 0.71 for H3K27ac, H3K4me3, and H3K27me3, respectively; **Supplementary Fig. 3**). To ensure that tagmentation reflected true histone modifications rather than intrinsic bias of Tn5 for open chromatin, we evaluated concordance across marks and against external datasets. The two activating marks showed increased correlation (Spearman’s ⍴ = 0.76; **Fig. 1c**), whereas the correlation between activating and repressive marks was substantially lower (Spearman’s ⍴ = 0.31 and 0.36 for H3K27me3 *vs.* H3K27ac and H3K4me3, respectively; **Fig. 1c**). Comparison with an external ATAC-seq dataset^26^ further supported signal specificity, with high chromatin accessibility observed at SPACE-Tag H3K27ac peaks and minimal ATAC-seq enrichment at SPACE-Tag H3K27me3 peaks (**Supplementary Fig. 4**). Together with strong concordance with ENCODE reference datasets (H3K27ac and H3K4me3 Spearman’s ⍴ > 0.88; **Fig. 1d**) and high reproducibility across biological replicates (Pearson’s r > 0.94; **Supplementary Fig. 5**), these results demonstrate that SPACE-Tag profiles the spatial epigenome with high fidelity.

Consistent with its sensitivity and specificity, SPACE-Tag also captured biologically meaningful spatial structure in intact tissues. Applying an unsupervised, spatially aware topic model^27^ (**Methods**) to the H3K27ac profiles resolved domains corresponding to the isocortex, piriform cortex, thalamus, hypothalamus, dorsal thalamus, fiber tracts, and the hippocampus (**Fig. 1e**). These regional topics recapitulate broad domains previously identified by DBiT-seq, further underscoring the sensitivity of SPACE-Tag to accurately map established brain anatomical structures (**Supplementary Fig. 6)**. Clustering based on H3K4me3 and H3K27me3 profiles yielded comparable results, but exhibited less spatial specificity (**Fig. 1e**). This pattern is consistent with the role of H3K27ac in marking cell-type-specific enhancer elements^28^. The activation and repression scores inferred from the histone modification profiles recapitulated established spatial gene expression patterns of canonical markers^29^ (**Fig. 1f,g, Supplementary Fig. 7).** For example, the neuronal transcription factor *Satb2*, which drives formation of cortical neuronal projections^30^, showed increased H3K27ac and H3K4me3 activation scores in the isocortex, while its corresponding H3K27me3 repression scores were reduced in this region. Consistent patterns were observed for *Tcf7l2* marking the thalamus^31^, and *Zeb2*, a marker of a joint region representing the isocortex, fiber tracts and the hippocampus. These inferred gene activity maps also closely mirrored spatial expression patterns captured by MERFISH^29^ (**Fig.1g**).

Finally, by integrating H3K27ac SPACE-Tag data with reference single cell profiles^32,33^ (**Methods**), we successfully deconvolved each spatial spot to infer cell type composition across the tissue (**Fig. 1h**). Confirming SPACE-Tag’s capacity to resolve biologically relevant spatial patterns, distinct cell types were consistently predicted in the expected anatomical regions: glutamatergic neurons were predominantly localized to the isocortex, oligodendrocytes to fiber tracts, and astrocyte subtypes segregated between the hippocampus and thalamus (**Fig. 1h**). Spatial analysis of active super-enhancers (SEs) and their associated TFs revealed three distinct modules that co-varied across tissue regions and cell types (**Supplementary Table 1, Supplementary Fig. 8, Methods**). Module 1, enriched for *Gata4*, localized primarily to excitatory and inhibitory neurons in the isocortex. Module 2 localized to the hypothalamus astrocytes, where it was linked to regulating axon regeneration and neuronal activity^34^. Finally, Module 3 localized to the oligodendrocytes in white matter tracts where we observed enrichment of *Mbd1* and *Srebf2*^35^, which are factors implicated in the maintenance of neural stem cell identity^36^ and astrocytic cholesterol metabolism^37^ (**Supplementary Fig. 8, Supplementary Table 1**). This correspondence between inferred cell types, gene activity patterns, functional elements and anatomical domains demonstrates that SPACE-Tag-derived histone modification profiles are sufficient to resolve spatial functional organization across intact tissues.

Having established that SPACE-Tag can map the spatial organization of chromatin states, cell types, and gene regulatory programs across intact tissues, we next applied this approach to aging mouse brain and generated a large-scale spatial epigenomics dataset across the adult life span (**Supplementary Fig. 9)**. Brain aging has often been characterized as global epigenetic erosion^38^, yet recent evidence revealed that it is a spatially heterogeneous process^39,40^. Because age-associated transitions involve coordinated remodeling of active and repressive chromatin states^41,42^, resolving these dynamics requires simultaneous spatial measurement of multiple histone modifications within intact tissue architecture. To systematically examine the spatial organization of epigenetic aging, we leveraged the scalability of SPACE-Tag and established a multimodal spatiotemporal atlas of the mouse brain across the adult lifespan (N_animals = 4, N_sections = 24, and N_spots = 95,442) encompassing young adult (3 months), middle age (12 months), late-middle age (18 months), and aged (24 months) timepoints (**Fig. 2a**). We profiled five key histone modifications marking different genetic elements: H3K27ac (promoter and enhancer), H3K4me1 (enhancer), H3K4me3 (promoter), H3K27me3 (repressive chromatin), and H3K9me3 (heterochromatin) together with the transcriptome. These modalities were profiled in consecutive sections, followed by computational registration and integration into a unified coordinate system (**Methods**).

**Figure 2.**
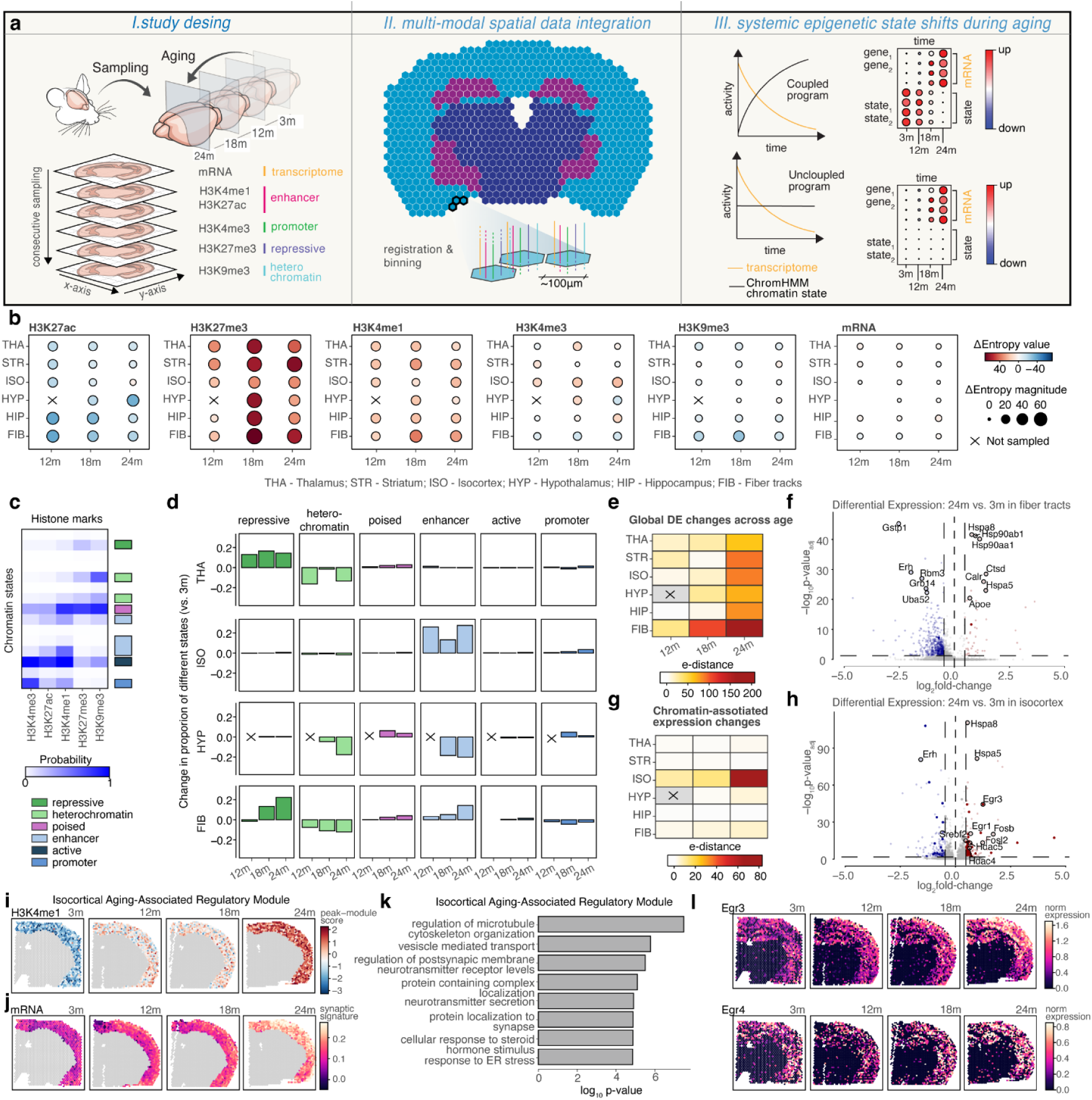
SPACE-Tag profiling of mouse brain aging. (**a**) Construction and analysis of a spatial multi-omic atlas of mouse brain aging. Left: Experimental workflow. Serial consecutive coronal mouse brain sections were collected at 4 time points and subjected to SPACE-Tag profiling of five histone modifications alongside spatial transcriptomics. Middle: image registration and binning were used to construct an integrated multimodal dataset with an effective spatial resolution of 100 μm. Right: Integrative analysis of histone and transcriptomic modalities to identify coupled and uncoupled programs of transcriptional and epigenetic shifts during aging. (**b**) Dotplots of global changes in tissue epigenetic entropy per tissue and histone mark compared to 3m timepoint. Color shows change in entropy value; dot size represents magnitude of change. (**c**) Heatmap of emission probabilities for 15 inferred chromatin states (rows) by chromHMM across 5 histone marks (columns). Color shows the probability of observing each histone mark (columns) at each inferred chromatin state (rows). Right annotation (color code) shows manual annotation of states based on histone probabilities. (**d**) Barplot of global changes in chromHMM chromatin states at differential peaks in selected tissue regions compared across aging as compared to 3m. (**e**) Heatmap of global changes in gene expression in aged tissue regions compared to 3m. Color represents magnitude of transcriptional change as measured by E-distance (**Methods**). (**f**) Volcano plot of differentially expressed genes in fiber tracts at 24m compared to 3m; color represents direction of expression change, red for upregulated genes and blue for downregulated. Solid points represent genes associated with proximal chromatin shifts; transparent points represent genes without proximal chromatin shifts. Dashed lines represent the significance thresholds for adjusted p-value (0.05) and fold change (0.5). Circled spots represent known transcription factors and chromatin remodellers. (**g**) Heatmap of gene expression changes associated with epigenetic shifts in aged tissue regions compared to 3m. Color represents magnitude of transcriptional change as measured by E-distance. (**h**) Volcano plot of differentially expressed genes in isocortex at 24m compared to 3m; color represents direction of expression change, red for upregulated genes and blue for downregulated. Solid points represent genes associated with proximal chromatin shifts; transparent points represent genes without proximal chromatin shifts. Dashed lines represent the significance thresholds for adjusted p-value and fold change. Circled spots represent known transcription factors and chromatin remodellers. (**i**) Spatial plots of isocortex H3K4me1 peak module scores across mouse brain aging. Color represents normalized module signal. (**j**) Spatial plots of synaptic gene signature across mouse brain aging. Color represents normalized signature expression. (**k**) Barplot of GO overrepresentation analysis of consensus isocortex gene signature, ranked by adjusted p-value. (**l**) Spatial gene expression plot of activity-dependent neuronal TFs *Egr3* and *Egr4* across mouse brain aging. Color represents normalized gene expression.

Leveraging epigenetic entropy as a measure of the degree of disorder across epigenetic states^43–45^ (**Methods**), we found that aging trajectories displayed significant heterogeneity across modalities and anatomical regions. While most modalities showed small, localized fluctuations across regions, H3K27me3 and H3K4me1 exhibited consistently increased entropy (**Fig. 2b**). The entropy shifts in chromatin states exceeded those in the transcriptome, indicating that the epigenome undergoes more substantial remodeling with age and may provide a more sensitive marker of cellular aging. To explore this further, we inferred combinatorial chromatin states using ChromHMM^46^ (**Methods**), and identified 15 distinct states defined by co-occurring histone marks, corresponding to enhancer, promoter, activating, poised, repressive and heterochromatin states (**Fig. 2c**). Quantifying age-associated transitions at differential peaks revealed that chromatin remodeling was primarily associated with shifts in repressive, heterochromatin, and enhancer states, and that anatomical regions exhibited distinct trajectories (**Fig. 2d; Supplementary Fig. 10**). For example, aging in the fiber tracts was characterized by expansion of repressive and enhancer states, driven primarily by accumulation of H3K27me3 and H3K4me1, respectively, alongside a depletion of H3K9me3-rich constitutive heterochromatin. In contrast, the hypothalamus showed a synchronized decline in both heterochromatin and enhancer states, evidenced by loss of H3K9me3 and H3K4me1. Finally, the isocortex followed a unique trajectory marked by a progressive, age-dependent gain in enhancer-associated states, associated with increased H3K4me1. Compared to the epigenome, transcriptome-wide analysis (**Methods**) revealed that all brain regions exhibited age-associated mRNA shifts (**Supplementary Table 2**), with fiber tracts showing the highest overall transcriptome-wide deviation at the aged timepoint (**Fig. 2e**). Consistent with previous reports^47^, these changes were dominated by widespread downregulation (**Fig. 2f**), affecting, among others, genes associated with metabolic function and homeostasis (*Gstp1*, *Grb14*, *Uba52*), as well as cell proliferation and survival (*Rbm3*, *Erh*). A smaller subset of upregulated genes was enriched for stress response pathways (*Hspa8*, *Hsp90aa1*, *Hsp90ab1*, *Hspa5*, *Calr*), inflammatory processes (*Ctsd*), as well as genes associated with neurodegenerative disease such as *Apoe*.

To investigate the extent to which these age-related transcriptional changes were accompanied by epigenetic remodeling, we stratified differentially expressed (DE) genes according to whether they exhibited concurrent local chromatin state transitions (**Methods**) (**Fig. 2g**). Strikingly, the relationship between transcriptional and chromatin changes differed substantially across brain regions. While most transcriptional changes in fiber tracts occurred without detectable chromatin state transitions at nearby regulatory regions, in the isocortex they were largely coupled to local chromatin remodeling (**Fig. 2g**). This was further supported by the significant differential expression of several transcription factors and chromatin remodelers (**Fig. 2h**). Consistent with this coupling, we identified a distinct module of regulatory regions in the isocortex that gained H3K4me1 signal with age (**Fig. 2i, Supplementary Table 3**). These regions overlapped with SEs more frequently than with regular enhancers (**Supplementary Fig. 11a, Methods**), indicating that age-related H3K4me1 increase in the isocortex is concentrated at high-occupancy regulatory hubs. Notably, this change occurred without a corresponding increase in H3K27ac (**Supplementary Fig. 11b**), suggesting selective enhancer remodeling rather than a global increase in enhancer activation. Consistent with this hypothesis, genes associated with this peak module were uniformly upregulated with age and were strongly enriched for synaptic signaling and neuronal function, including key regulators of neuronal plasticity such as Egr3 and Egr4 (**Fig. 2i-l**). In addition, motif enrichment and transcriptional analyses converged on the EGR family transcription factors, identifying Egr1 and Egr3 as both enriched at these regions and transcriptionally upregulated with age (**Supplementary Fig. 12**). To evaluate this further, we examined independent single-cell transcriptomic datasets of mouse cortical development^48^ and found this signature associated with postnatal synaptic pruning in glutamatergic neurons (**Supplementary Fig. 13**). In contrast, analysis of a spatial transcriptomic dataset from an Alzheimer’s disease mouse model^49^ revealed downregulation of this same gene set in disease (**Supplementary Fig. 14**). We therefore hypothesize that this observed age-dependent H3K4me1 expansion is driven by an EGR-mediated recruitment of chromatin-modifying complexes to high-occupancy regulatory hubs. This model is supported by a documented molecular hierarchy, wherein EGR-family proteins physically interact with the co-activators p300/CBP^50^, which in turn scaffold the MLL4/KMT2D complex^51^, the primary methyltransferase responsible for depositing H3K4me1. The functional importance of this interaction is further supported by previous findings showing that *Egr3* deficiency leads to a marked depletion of H3K4me1 at its genomic targets^52^. Together, these findings point towards an EGR-MLL4-driven enhancer remodeling in the aging isocortex potentially reflecting an adaptive regulatory program that supports neuronal function during healthy aging.

In contrast to methods built on custom microfluidic architectures, which require specialized fabrication and infrastructure, SPACE-Tag leverages the widely adopted^53^ and commercially accessible Visium platform, enabling direct integration of CUT&Tag profiling into established spatial transcriptomic workflows. Combined with T7-based linear amplification to overcome signal sparsity, SPACE-Tag uniquely enabled the detection of region-specific epigenetic rewiring, revealing spatially structured aging trajectories that would remain obscured in bulk or single-modality approaches. While the current implementation is limited by spot-level spatial resolution and primarily focuses on histone modifications, future developments are expected to move toward single-cell resolution and simultaneous multi-modal profiling.

## Materials and Methods

### Murine tissue collection

C57BL/6J mice were obtained from The Jackson Laboratory (Bar Harbor, ME) and maintained following ethical guidelines monitored by the Institutional Animal Care and Use Committees (IACUC) established by the Division of Comparative Medicine at Columbia University, under protocol AACE0752. All mice were maintained under SPF conditions on a 12-h light–dark cycle at an ambient temperature of 21.5 ± 1 °C with relative humidity between 30% and 70% and were provided food and water ad libitum. Brains were collected immediately after cervical dislocation and washed with ice-cold 1X PBS. Tissues were dried and embedded in Optimal Cutting Temperature (23-730-571, Fisher Scientific) in 22×22×20 mm disposable molds (15160-215, VWR). Samples were then placed onto a pre-chilled metal plate and put on dry ice for 2 min or until completely frozen. Samples were transferred and stored at -80°C until use.

### Immunostaining and epifluorescence microscopy

To optimize histone antibody concentrations, we performed the following experiments. Frozen 10 μm tissue sections were mounted on microscopic slides and incubated at 37°C for 5 min on an Eppendorf ThermoMixer C with a thermal block. After fixation in 0.2% formaldehyde (F8775, Sigma Aldrich) at room temperature (r.t.) for 10 min, tissue sections were quenched with 1.25 M Glycine in 1X PBS, followed by a 1X PBS wash. Tissue sections were permeabilized in 1X Perm/Wash buffer (554723, BD) for 30 min at 4°C. Antibodies were added at desired dilutions in 1X Perm/Wash buffer and incubated for 1 h at r.t. Subsequently, tissues were washed with 1X PBS and incubated for 30 min with fluorescently labeled secondary antibody (711-165-152, Jackson ImmunoResearch) in CT wash buffer (20 mM HEPES, 150 mM NaCl, 0.5 mM spermidine, proteinase inhibitors, and 0.01% digitonin). Stained tissues were then washed for 5 min in100 μl CT wash buffer and then proceeded to microscopy. Samples were imaged using a Zeiss Axioscope inverted microscope with a 20× objective (MetaSystems). Images were stitched using Vslide (v1.1.132) and extracted with VSViewer in a JPEG format. The following antibodies were optimized and used in spatial CUT&Tag experiments: anti-H3K4me3 (ab213224, Abcam) anti-H3K27me3 (9733S, Cell Signaling), anti-H3K27ac (ab4729, Abcam), anti-H3K4me1 (ab8895, Abcam) and anti-H3K9me3 (ab8898, Abcam).

### Pseudo-spatial slide production

Amino-binding CodeLink high-density (HD) glass slides (DHD1-0023, Surmodics) were printed with an equimolar mix of four amine-modified oligonucleotides containing a poly(dT) segment (Amine-visium2-1/2/3/4, **Supplementary Table 4**), following an established protocol ^54^. Briefly, slides were placed in 16-well hybridization cassettes (AHC4X16, ArrayIt), and a printing solution (150 mM Sodium Phosphate, 0.06% sarkosyl, pH 8.5) containing 20 µM of amine-modified oligonucleotides (**Supplementary Table 4**) was added to each well and incubated for 1 min. After removing the solution, slides were air-dried at room temperature for 5 h, followed by a 14 h incubation in a high (>80%) humidity chamber at r.t. The slides were then blocked for 30 min in a blocking solution (50 mM Tris, 50 mM Tris-HCl (pH 9), and 50 mM ethanolamine) at 50 °C, followed by a 30 min wash in 4X SSC with 0.1% SDS at 50 °C. Finally, the slides were washed in Milli-Q water, spin-dried, and stored in a desiccator (RH<10%) until use. These slides were used in all optimization experiments (**Supplementary Fig. 3**).

### Transposome assembly

Firstly, each modified Tn5 adaptor (T7L_MedsB or dual_ME19_polyT_v2, 25 μl of 100 μM) was mixed separately with 25μl mosaic ends oligonucleotides (ME_19_Phos, 100 μM) and 1μl 50X annealing buffer (0.5 M Tris pH 8, 2.5 M NaCl). The mixtures were annealed by heating to 95°C for 5 min, followed by gradual cooling to 14 °C at -0.1 °C/s. ProteinA-Tn5 was diluted to 0.5 mg/ml in dilution buffer (50% glycerol, 10mM Tris-HCl, pH 7.5, 100 mM NaCl, 0.1 mM EDTA, and 1 mM DTT) and then combined with the annealed adaptor at a 5:1 ratio. The reaction was incubated at r.t. for 1 h to form the transposome. The transposome was stored at -20 °C until use. All oligonucleotide sequences are available in **Supplementary Table 4**.

### SPACE-Tag procedure

#### Tissue fixation, permeabilization, and antibody staining

Frozen tissue sections were prepared using a Leica CM1950 Cryostat at -17°C and sectioned at a thickness of 10 μm to be mounted onto capture areas on Visium slides (1000184, 10x Genomics). The slides were then incubated at 37°C on an Eppendorf ThermoMixer C with a thermal block for 5 min to enhance tissue adhesion. For fixation, tissue sections were treated with 1 mL 0.2% formaldehyde (F8775, Sigma Aldrich) in 1X PBS for 10 min, followed by quenching with 1.25 M glycine in PBS for 5 min. Sections were then washed with 1X PBS for 2 min. Visium slides were then mounted on ArrayIt reaction cassettes for the following experiment (AHC1X16, ArrayIt). Permeabilization was performed with 70 μl 1X BD Perm/Wash buffer (554723, BD) for 30 min at 4°C. Primary antibodies were diluted 1:50 (9.8–20.0 µg/ml) in 70 μl 1X BD Perm/Wash buffer and incubated for 1 h at r.t. Tissue sections were then washed with 1X PBS for 5 min r.t. and incubated with guinea pig anti-rabbit IgG secondary antibody (NBP1-72763, Novus Biologicals; 1:50, 24 µg/ml) in 70 μl CT wash buffer (20 mM HEPES, 150 mM NaCl, 0.5 mM spermidine, proteinase inhibitors, and 0.01% digitonin) for 30 min at r.t.

#### pA-Tn5 Tethering and tagmentation

Per capture area, antibody-stained sections were incubated with preloaded ProteinA-Tn5 (MacroLab, Berkeley) at a concentration of 0.01 mg/ml in 70 μl 300-wash buffer (20 mM HEPES, 300 mM NaCl, 0.5 mM spermidine, proteinase inhibitors, and 0.01% digitonin) at r.t. for 1 h. Unbound ProteinA-Tn5 was then removed by washing with 100 μl of the 300-wash buffer for 5 min. Following this, tagmentation was initiated by adding 70 μl of tagmentation buffer (10 mM MgCl_2_ in 300-wash buffer) and incubated at 55°C for 1 hour. Alternatively, we tested tagmentation at 37°C for 1 hour, and these results are presented in **Supplementary Fig. 2**. Tagmentation was stopped by adding 70 μl 0.1 M HCl (H9892, Sigma-Aldrich) for 2 min at r.t., which also served as the acid-clearing step, followed by a wash with 1X NEBuffer 2 (B7002S, NEB).

#### Gap filling and in vitro transcription

Gap filling extends the genomic 3′ ends across the transposition gaps and the single-stranded adaptor overhangs, making the T7 promoter double-stranded for transcription. Per capture area, the gap-filling reaction was performed on the tissue using a 70 μl reaction mixture containing 1X NEBuffer 2 (B7002S, NEB), 0.5 mM dNTPs (N0447S, NEB), 28 U RNaseOUT (10777019, Thermo Fisher Scientific), and 10 U exo-Klenow Fragment (M0212L, NEB). The reaction was incubated at 37°C for 30 min and removed. The sections were washed with 1X transcription buffer (EP0111, Thermo Fisher Scientific) for 5 min at r.t. *In vitro* transcription (IVT) was initiated by adding 70 μl IVT reaction mixture containing 1X transcription buffer (EP0111, Thermo Fisher Scientific), 4 mM of each rNTP (N0466S, NEB), 16 mM additional MgCl₂ (20303M, Sigma), 140 U RNaseOUT (10777019, Thermo Fisher Scientific), and 14 U T7 RNA Polymerase (EP0111, Thermo Fisher Scientific) to the tissue sections. The reaction was incubated overnight (∼14 hours) at 37°C. We alternatively tested T4-based gap filling using 1X T4 ligase buffer, 0.05 mM dNTPs, 2 U/μL T4 DNA ligase (M0202S, NEB), and 0.015 U/μL T4 DNA polymerase (M0203S, NEB) in 70 μl reaction volume at r.t for 30 min. These comparison results are presented in **Supplementary Fig. 2**.

#### H&E staining

After removing the IVT mixture, tissue sections were removed from the reaction cassette and fixed in methanol at -20°C for 30 min, and an abbreviated hematoxylin and eosin protocol was used for histological staining. Briefly, tissue sections were incubated in isopropanol for 1 min and in Mayer’s hematoxylin (S330930, Agilent) for 7 min. Sections were then rinsed by immersing the slides in deionized water. Subsequently, the slides were incubated for 2 min in Bluing Buffer (CS70230, Agilent) and rinsed in deionized water. The slides were then incubated for 1 min in Eosin Y solution (100 μl Eosin Y (HT110216, Sigma Aldrich) in 900 μl 0.45M Tris-acetate, pH 6) and rinsed in deionized water. Finally, the slides were dried and imaged. Tissues were imaged using a Zeiss Axioscope microscope with a 20× objective. Images were stitched with Vslide (v1.1.132) and extracted with VSViewer in a JPEG format.

#### Reverse transcription (library preparation)

After imaging, the slides were mounted again on the reaction cassettes and washed with 1X Maxima H Minus Reverse Transcriptase buffer at 37°C for 30 min. The buffer was then removed, and a reverse transcription (RT) mixture was prepared according to the Visium protocol with slight modifications. Briefly, 43.06 μL H_2_O, 18.75 μL RT reagent, 1.5 μL reducing reagent B, and 11.69 μL RT enzyme D are mixed for each well of reaction. After adding the RT mixture, tissue sections were incubated at 53°C for 1.5 h for cDNA synthesis.

#### Reverse transcription (fluorescent cDNA gene activity print)

After imaging, the slides were remounted on the reaction cassettes and washed with 1X Maxima H Minus Reverse Transcriptase buffer at 37°C for 30 min. The buffer was then removed, and a 70 μL reverse transcription (RT) mixture was prepared as follows: 39.6 μL Water, 7 μL 10xRT buffer (B0253S, NEB), 0.65 μL BSA (20 mg/ml), 3.5 μL QC dNTPs (10mM dATP, 10 mM dGTP, 10 mM dTTP, 0.25 mM dCTP), 1.75 μL cy3-dCTP (0.2 mM, PA53021, Cytiva), 3.5 μL RNaseOUT (10777019, Thermo Fisher Scientific), 7 μL M-MuLV Reverse Transcriptase (M0253S, NEB). The RT mixture was added to the tissue sections, which were incubated at 37°C overnight for cDNA synthesis. Next day, the RT mix was removed and 70 μL proteinase K mixture was added per well (9 μL proteinase K [EO0491, Thermo Fisher Scientific] and 61 μL PKD buffer [1034963, Qiagen]). Tissue removal was performed by incubating 55 °C for 1hr. Slides were then washed sequentially in 2xSSC with 0.1% SDS at 50 °C for 10 min, 0.2X SSC at r.t. for 1 min and 0.1X SSC at r.t. for 1 min. Finally, the slides were imaged on a Dragonfly 600 confocal microscope system (Andor, Belfast, UK) to capture the cDNA print, with image acquisition performed in Fusion software (v2.4.0.14). Imaging was carried out using a 594 nm laser at 20% intensity with an exposure time of 200 ms. TIFF images were analyzed using ImageJ. For each sample, three regions of interest (ROIs) were selected from background areas to measure background fluorescence intensity, and three additional ROIs were selected within the tissue cDNA print area. Within the tissue, ROIs were placed in the hippocampus, isocortex, and hypothalamus.

#### Second-strand synthesis

The RT mixture was removed, and 70 μl 0.08 M KOH (freshly diluted from the 8 M KOH stock, 11-102-0056, Thermo Fisher Scientific) was added to the tissue sections and incubated for 5 min at r.t. to remove the RNA strand. KOH was then removed, and Elution Buffer (EB; 10 mM Tris-Cl, pH 8.5) was added to wash the tissue sections. A primer annealing mix of 70 μl containing 2× SSC, 20% formamide (221198, Sigma-Aldrich), 0.1% Tween-20, and 0.02 µM PF-nextera-R2 (sequences available in **Supplementary Table 4**) was applied to the tissue sections and incubated at r.t. for 30 min. The unbound primer was washed away with EB. Second-strand cDNA synthesis was performed using 70 µl second-strand mix containing 5 U/μl Klenow Fragment (M0210L, NEB), 1 µM PF-nextera-R2 primer (sequences available in **Supplementary Table 4**), 1mM dNTP mix, and 1X Maxima H Minus Reverse Transcriptase buffer. The reactions were incubated at 37°C for 1 hour. Following the reaction, tissue sections were first washed with EB. Then, 35 μL 0.08 M KOH is added to the tissue, incubated for 10 min at r.t., and the solution volume collected. 5 μL of Tris buffer (1 M, pH 7.0, BBT-70, Boston Biosciences) was added to each sample to neutralize the KOH.

#### Library preparation and indexing

PCR amplification was performed by adding PCR mix and primers to the neutralized sample to reach a concentration of 1X NEBNext High-Fidelity PCR Master Mix, 0.5 µM cDNA Oligo 1 primer, and 0.5 µM PF-NexteraR2 (oligonucleotide sequences available in **Supplementary Table 4**). After amplification, libraries were cleaned using a 0.65:1 ratio with AMPure XP beads (A63880, Beckman) and eluted in 40 μL of water. Indexing PCR was performed using 10 μL of the PCR product in 1X NEBNext High-Fidelity PCR Master Mix and 0.5 µM indexing primers (SIPCR-T501-T504, N701-N722, oligonucleotide sequences available in **Supplementary Table 4**). Indexed cDNA libraries were then purified using AMPure XP beads at a 0.6:1 ratio, following the manufacturer’s protocol, and eluted in 40 µL of EB. All samples were sequenced on the Illumina Novaseq X sequencing platform using a 28-cycle forward read (R1), 8-cycle index read, and 100-cycle reverse read (R2) configuration (final loading concentration of 190 pM).

### Data preprocessing and statistical quantification

#### Primary data processing

A custom computational workflow was implemented using the Snakemake workflow manager v7.32.4 to perform data preprocessing. Reads were first trimmed with cutadapt v4.5^55^. Spatial barcode sequences and UMIs are next extracted from the forward read (R1), and barcode filtering was performed using the 10X Visium V1 barcode list as a reference with umi-tools v1.1.4^56^. The reverse read (R2) was then aligned using local alignment with bowtie2 v2.5.2^57^ to the mm10 mouse genome reference. Aligned sam files are deduplicated using the spatial barcode and alignment coordinates before peak calling using macs2 v2.2.9.1^58^ with custom parameters ‘--slocal 5000 --local 50000 --broad --max-gap 1000’ to enable identification of broad histone modification domains. The resulting peaks list and alignment files were used to create a fragments file and raw peak-by-spot data matrix using sinto v0.10.0 (https://github.com/timoast/sinto). The raw peak matrix was next filtered to remove peaks associated with polyadenylated genome sequences. A custom script was implemented in R v4.3.1 to identify peaks associated with poly-A and poly-T sequences of length 20 within 70 bp of the peak, and these peaks are filtered out. In addition, the tissue region was annotated using manual fiducial alignment using the Loupe browser v6.3.0 (10x Genomics), and spots outside the tissue were filtered out. We further manually annotated H&E images of the mouse brain using a custom in-house interactive tool^16^, using annotation labels corresponding to the retrosplenial area, visual cortex, somatosensory cortex, auditory cortex, perirhinal area, piriform cortex, fiber tracts, hippocampus, thalamus, hypothalamus, striatum, and amygdala. The resulting filtered peak-by-tissue spot matrix was used for downstream analysis.

#### Comparison of estimated library complexity across conditions

To compare differences in library complexity between conditions for protocol optimization (**Supplementary Fig. 2**), we used Preseq^59^, an empirical Bayesian method for estimating the total number of unique fragments in a cDNA library conditioned on the observed frequency of duplicated fragments. Specifically, we used the ‘bound_pop’ method to compute the total library complexity estimate.

#### Tissue registration

To register mouse tissue sections, we leveraged manual spot annotations together with the highly stereotyped architecture of the brain. This annotation-guided strategy provides greater robustness than direct alignment of H&E images, which are often confused by incomplete or disrupted morphologies. Our registration workflow proceeds as follows. First, for each replicate, we identify centroids corresponding to annotated spots in the retrosplenial area and thalamus. We then rotate the spatial coordinates of spots within each replicate to minimize the distance between corresponding centroids across replicates. After rotation, we translate the coordinates so that the point equidistant between the centroids lies at the origin of the coordinate system. Finally, all registered sections are manually inspected to confirm alignment accuracy.

#### Gene activity quantification

The ‘GeneActivity’ function in Signac v1.13.0^60^ was used to estimate gene activities from histone modification levels. To that end, reads aligning in the gene body and within 2 kb upstream of the gene body were aggregated to give a gene activity score for each gene. Joint quantification of gene activity across histone modifications and experimental replicates was then performed using Splotch v0.1.0, a hierarchical Bayesian model for spatial counts data^61^. Splotch models counts data across multiple spatial experiments using a hierarchical model, wherein the transcript count for each gene (or, in our case, gene activity of each gene) *i* in spot *k* for sample *j* is assumed to be sampled from a Poisson distribution. The model specification is given in Eq. 1 below:

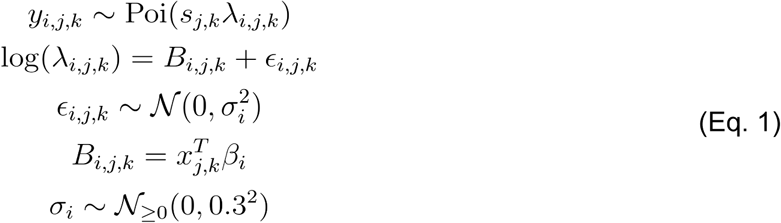

Here, ∈*_i,j,k_* is an error term to account for spot-level variability, and *S_j,k_* is a size factor, corresponding to the total UMIs in spot *i* (*i.e.* 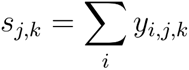), to normalize for per-spot read depth. A two-level hierarchical prior is placed on the latent rate parameter ⋋ to account for differences between animals and distinct histone modifications (Eq. 2). Specifically, this is done through β*_i_*, which quantifies the contribution of each annotation level in the one-hot encoded annotation vector 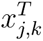 to the expression of gene *i* in spot *k*, for sample *j*:

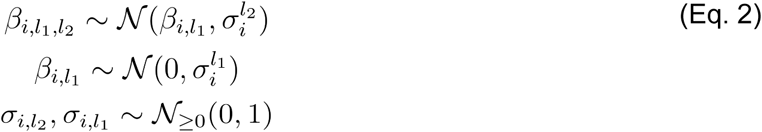

Here, we set the upper-level *l*_1_ annotation to be the histone modification identity, and the *l*_2_ annotation to the identity of the experimental animal. Model inference was performed using Markov Chain Monte Carlo sampling with Stan v2.36.0 (https://mc-stan.org), with 4 chains of 500 burn-in and 500 sampling iterations each. To plot inferred gene activity across replicates, registered coordinates (see above) were used. The mean gene activity of each spot was computed using the inferred posterior means of ⋋ and plotted.

### Comparisons to published data

To establish the sensitivity and specificity of SPACE-Tag, we performed comparisons against a variety of bulk, single-cell, and spatial datasets.

#### Comparison with bulk ChIP-seq

To validate the fidelity of SPACE-Tag relative to established benchmarks, we compared our spatial data with adult mouse isocortex H3K27ac and H3K4me3 ChIP-seq data from the ENCODE portal (H3K4me3: ENCSR943LBN; H3K27ac: ENCSR491RBV). We first generated pseudo-bulk profiles by aggregating SPACE-Tag signals from spots assigned to specific anatomical domains via spatially-aware probabilistic topic modeling (STAMP v0.1.0)^27^. Raw counts were aggregated across these spots to create a combined topic profile. For the reference, we utilized peak-level counts from ENCODE aligned BAM files. To account for differences in sequencing depth and library size, both the SPACE-Tag pseudo-bulk and ENCODE reference profiles were normalized to Counts Per Million (CPM). To quantify the concordance between the two platforms, we calculated pairwise Spearman correlation coefficients rho on the log-transformed log_10_(CPM + 1) values. To ensure a robust comparison of shared regulatory features, we excluded zero-count peaks and performed density-based visualization using the MASS::kde2d function (v7.3.65).

#### Benchmark SPACE-Tag Specificity to ATAC-seq data

To evaluate the spatial fidelity and biochemical specificity of SPACE-Tag, we compared our profiles against an adult mouse prefrontal cortex ATAC-seq^26^ (GEO: GSM3034633). For this comparison, SPACE-Tag BAM files were generated by aggregating signals from spots assigned to cortical spatial domains, as identified via spatially-aware probabilistic topic modeling (STAMP v0.1.0). The resulting cortical-specific BAM files were converted to BigWig format with 1x genome coverage normalization using deepTools (v3.5.5)^62^. To assess the functional accuracy of our mapped epitopes, we utilized deepTools (v3.5.5)^62^ to center the bulk ATAC-seq signal on high-confidence peaks identified in our spatial data (±3kb window). Using the computeMatrix and plotHeatmap modules, we visualized the distribution of accessibility across these regions to confirm the biochemical specificity of our spatial profiling.

#### Comparisons with spatial CUT&Tag

To compare with existing spatial CUT&Tag methods, we downloaded processed fragments files from the GEO repositories for the Deng *et al*., Zhang *et al*., and Guo *et al*. studies^10,11,63^ (GSE165217, GSE205055, and GSE263333, respectively); we then computed the number of fragments per barcode using the fragments file.. We compared the total fragment count and FRiP score between these methods and our SPACE-Tag data. For joint integrative analysis between SPACE-Tag and CUT&Tag, we identified a spatial CUT&Tag dataset (GSM6704979) of mouse brain H3K27ac, requantified the fragments file using peaks called on SPACE-Tag samples, then performed joint topic modelling as described below.

#### Comparison with single-cell CUT&Tag

To compare with existing single-cell CUT&Tag methods, we used the publicly available preprocessed peak-by-cell matrix (accessed from the GEO repository GSE198467). We first merged this data with SPACE-Tag data by re-quantifying the counts matrix of SPACE-Tag data using peaks identified in the scCUT&Tag dataset. Next, we performed anchor-based integration with Seurat v5.2.1^60^ for our SPACE-Tag data with ‘FindTransferAnchors’ and ‘TransferData’ functions using the joint Latent Semantic Indexing (LSI) embedding of the merged data to compute neighbourhood distances, and plotted the resulting projection scores for the high-resolution cell subtypes identified in the original manuscript.

#### Comparison to Allen Brain Cell Atlas MERFISH

To compare cell type distributions between our inferred cell type projection scores derived from integration with single-cell CUT&Tag as described above with the annotated cell types from the Allen Brain Cell Atlas (ABC) MERFISH dataset^29^, we downloaded processed data from ABC version 20250531. We manually inspected serial sections along the anterior-posterior axis; this identified section MERFISH-C57BL6J-638850 to be visually similar in terms of overall morphology to our coronal sections, and we extracted the annotated cell subtype labels provided by the authors for this section.

### Analysis of gene regulation in the postnatal mouse brain

#### Topic modelling and clustering

To identify spatial domains in SPACE-Tag profiling, we performed topic modelling with STAMP v0.1.0, a spatially-aware probabilistic topic model^27^. Briefly, a joint topic model with n=10 topics was constructed with the registered data (see above) of all replicates of the adult brain, using the peak-by-spot count matrix with Poisson likelihoods for peak counts, and Simplified Graph Convolutions (option ‘sgc’) to account for the spatial locations of spots for inference of spatial domains. To ensure a consistent feature set between replicates, the union of peak sets across all replicates was used. To account for batch effects, the sample ID was included in the model as a categorical covariate. Training was performed for 2000 epochs with a learning rate of 0.01. To assign the resulting probabilistic topic proportions to discrete clusters, we filtered out topics associated with non-spatial technical variation, and labelled each spot using the topic with the highest proportion in that spot.

#### Analysis of cortical plate axis

To investigate functional region-specific H3K27ac peaks in the cortex of the P28 postnatal mouse brain, we first manually annotated functional regions across the brain and extracted functional regions corresponding to the cortex. We next defined a spatial cortical plate axis along the dorsal-ventral axis of the cortex. To do this, we first identified a set of anchor points in the tissue using K-means clustering (K=10), and then defined the axis by fitting a principal curve for the anchor point set. We then performed spatial module analysis on all peaks with greater than 100 counts across all spots, using Hotspot v0.9.1^64^, which estimates local correlations between spots given a pre-defined measure of similarity between spots. For cortical subregion analysis, the projection of each spot along the cortical plate principal curve was used as the similarity measure, from which a KNN neighbourhood graph (K=300) was constructed. The resulting peak-peak correlation matrix was then clustered using hierarchical clustering to obtain spatial modules.

#### Identification and analysis of super-enhancer elements

To identify putative super-enhancer elements, we adapted the strategy used in the original Whyte *et al*. publication^65^, by ranking H3K27ac and H3K4me3 peaks based on their aggregated normalized signal. The standard normalization method used in analysis of single-cell and spatial epigenomic data is the term frequency-inverse document frequency (TF-IDF) transformation, which is unsuitable for our purposes as it weights peaks by their inverse frequency; we therefore used an alternate normalization scheme whereby the raw fragment counts for each peak in a spot is normalized by a scaling factor equal to the ratio of the total number of unique fragments in that spot to the median unique fragments across all spots. This accounts for biases caused by sequencing depth and the number of nuclei per spot. Super-enhancer elements were then defined as peaks in the top decile of normalized signal that overlapped with an element in the SEdb database^66^. Motif enrichment in spatial modules was computed by comparing the number of motifs for TFs in the Cis-BP database^67^ in peaks associated with a spatial module versus a background set of 40000 GC-matched regions across the genome using the ‘FindMotifs’ function in Signac.

### Construction and analysis of a multimodal SPACE-Tag atlas of mouse brain aging

#### Image registration and sample integration

To construct an integrated multimodal dataset of transcriptomics and epigenetic modalities, we first independently preprocessed the data collected for each profiled histone modification as well as the gene expression modality to obtain filtered peak-by-tissue spot matrix as described above. We next performed image registration using ELD^68^. Spatial landmarks were manually defined using the H&E image across sequential sections representing distinct modalities using n=6 landmarks per section. The thin-plate spline function from ELD (ELD.warp.TPS) was then used with default parameters to perform locally non-linear registration of target images (corresponding to the epigenetic modalities) to the source image (corresponding to the transcriptomic modality). The resulting learned transform was then applied to spot coordinates to generate a common coordinate framework across Visium and multimodal SPACE-Tag spots. The aligned spots were then hexagonally binned using matplotlib.pyplot.hexbin from matplotlib v3.10.7, with gridsize=50 and mincnt=6, to produce a 50×50 grid of hexagonal bins. The resulting bins were filtered to those containing at least one spot from each histone and transcriptome modality. SPACE-Tag profiled fragments and Visium mRNA read counts were aggregated within each bin, and the resulting multimodal hexbin-by-feature matrix was used in all subsequent downstream analyses. Manual annotation was used to annotate brain tissue regions from H&E staining of the Visium transcriptomic data as described above; the resulting annotations were applied to hexbins.

#### Characterization of global trajectories of epigenetic and transcriptomic entropy

After generating annotated, binned multimodal spatial data, we next turned to characterising tissue annotation-specific changes in global epigenetic and transcriptomic state by assessing per-tissue entropy. Specifically, we wished to infer the entropy of the per-tissue hexbin distribution for each modality. Entropy estimation is difficult for high-dimensional distributions, such as the raw data matrices produced after multimodal integration; we therefore combined standard single-cell and spatial omics workflows, which involve feature selection and linear dimensionality reduction followed by construction of a k-nearest neighbor (KNN) graph in the low-dimensional embedding^69^, with nonparametric methods for entropy estimation using KNN graphs^70^. Specifically, we first constructed the KNN graph separately for each histone modality and the transcriptomic modality. For the histone modalities, we first binned the data into 10 kb genomic bins, before computing the term frequency-inverse document frequency transform using ‘RunTFIDF’ in Signac (v1.13.053). Next, we performed SVD using the ‘RunSVD’ function in Signac, took the top 50 components, and constructed a KNN graph (K=30). For the transcriptomic modality, we used log1p normalisation as implemented in ‘NormalizeData’ in Seurat, performed PCA on the top 2000 most highly variable features, then constructed a KNN graph (K=30) from the top 50 PCA components. For each KNN graph constructed per modality, tissue annotation and timepoint, we then estimated the entropy using the ‘entknn’ function from the R package copent v0.5.0^71^. Specifically, this estimates the Shannon entropy *H*(*X*) using the KNN estimator:

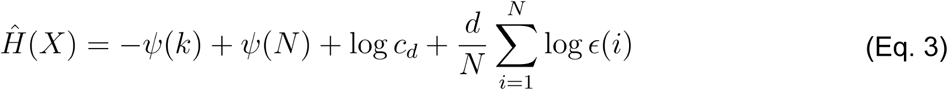

where *k* is the number of neighbours in the KNN graph, *N* is the number of points on the graph, *d* is the dimensionality of *X*, *c_d_* is the volume of a *d–*dimensional unit ball, and ϵ(*i*) is twice the distance between *x_i_* and its *k*th neighbour. Ψ(*x*) denotes the digamma function.

#### Chromatin state inference and analysis

We performed chromatin state inference using ChromHMM v1.27^46^. We first generated pseudobulk profiles of histone modifications across each tissue annotation at each timepoint by aggregating fragments across annotated hexbins. These profiles were binarized with 500bp resolution using the BinarizeBam function in ChromHMM. This resulted in 24 (4 timepoints by 6 tissue annotations) pseudobulk profiles each consisting of binarized signals in 500bp windows across 5 histone modifications. We then trained a HMM with k=15 hidden states on these pseudobulked profiles using the LearnModel function in ChromHMM, with n=5000 iterations. Inspecting the per-histone emission probabilities of this model, we observed a number of states associated with potential assay noise at polyadenylated genomic sequences, with high signal across all histone marks in all tissues and timepoints, and in addition were associated with the mm10 ENCODE blacklist^72^. We decided to filter out reads at genomic regions annotated as these ‘noise-associated’ states in greater than 15 of 24 pseudobulk profiles. We reran the ChromHMM state inference as described above on the filtered pseudobulk profiles, and used these filtered fragments files for all subsequent analysis.

We next quantified chromatin state transitions at genomic regions exhibiting differential histone modifications. To do so, we first performed differential peak analysis to find peaks with increased or decreased histone modification levels at later time points compared to 3m (3 months). This was done using the ‘FindMarkers’ function in Seurat, comparing the signal at each peak for each histone in each tissue region at 12m (12 months), 18m (18 months) and 24m (24 months) relative to 3m. We set the significance threshold at adjusted p.value < 0.05, with the additional constraint that the fraction of hexbins with non-zero signal at the peak must be above 0.3 in at least one of the two conditions in order to mitigate negative errors arising from signal sparsity. We then aggregated the resulting differential peak regions across all histones and time points for each tissue annotation to generate a union set of peaks exhibiting differential chromatin signal with aging in each tissue. We extracted the inferred chromatin states in this union peak set, and computed the proportion of each annotated chromatin state as a fraction of the total genomic width of the union peak set; we then calculated the difference in this proportion between each subsequent timepoint and the 3m timepoint to estimate the change in chromatin state proportion.

#### Analysis of coupled and uncoupled changes in the epigenome and transcriptome

We estimated the magnitude of global transcriptional changes associated with aging using E-distance, a general measure of distance between the distributions of single-cell expression datasets^73^. The E-distance *E(X,Y)* between two distributions is defined as:

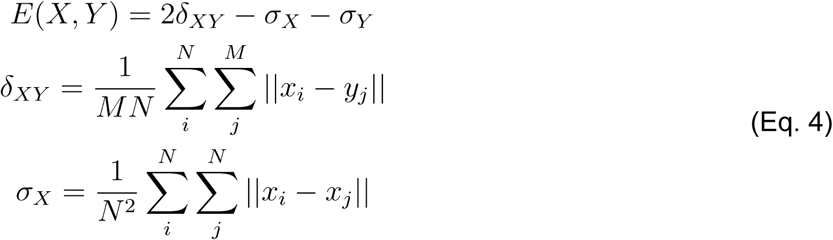

Where δ*_XY_* is the mean between-distribution and σ*_x_* the mean within-distribution squared euclidean distance between points. Following the original scPerturb^73^ workflow, we use the low-dimensional linear embeddings after normalization, feature selection and PCA as described above to generate the per-timepoint, per-tissue distributions *X,Y*. To estimate the magnitude of transcriptional changes associated with chromatin changes and thereby establish the degree of coupling between transcriptional and epigenetic trajectories within each tissue, we repeated this analysis, subsetting the raw hexbin-by-gene data matrix to the subset of genes that were within 10 kb of a differential peak from the union peak sets defined above. We performed normalization, feature selection and linear dimensionality reduction via PCA on these filtered genes, and recomputed E-distance using the distributions in principal component space induced by the filtered data matrix. In addition, we performed differential expression analysis within each annotated tissue region between the 24m and 3m timepoints using Seurat’s ‘FindMarkers’ function, setting the significance threshold at adjusted p.value < 0.05 and absolute log fold change > 0.5.

#### Analysis of cortex-specific changes in the epigenome and transcriptome

We first defined an isocortex-specific peak module by identifying differentially abundant H3K4me1 peaks at 24m compared to 3m in the isocortex, using a significance threshold of adjusted p.value < 0.05. We computed a chromatin module score using the ‘AddChromatinModule’ function in Signac, which calculates a z-score of the signal deviation at a given peak set against a set of matched background peaks with similar average signal and sequence characteristics. To investigate the association between this isocortex peak module and super-enhancer elements, we used our H3K27ac data to identify super-enhancers in conjunction with the SEdb database as described above. We then compared the proportion of super-enhancer and non-superenhancer elements, as defined by H3K27ac peaks, that overlapped with the isocortex peak module. Using our isocortex-specific peak module, we also constructed a consensus gene signature of genes that were both significantly differentially expressed in the isocortex at 24m compared to 3m, as well as within 10 kb of a peak in the isocortex peak module. We computed the gene expression signature score using the ‘AddModuleScore’ function in Seurat. In addition, we performed GO enrichment analysis for this signature using the ‘enrichGO’ function in clusterProfiler, as described above.

#### Comparing consensus isocortex gene signature with published data

We cross-referenced our consensus isocortex gene signature with a number of published datasets. For the single-nucleus mouse visual cortex developmental dataset in Gao *et al.*^48^, we grouped the cells using cell-type annotations provided by the authors, computed the gene expression signature score of our consensus gene signature using the ‘AddModuleScore’ function in Seurat, and performed linear regression of the signature score as a function of the number of days post-fertilization, assuming a 19-day gestational period. For the Visium cortex AD dataset^49^, we merged the data across all profiled mouse coronal brain sections, and jointly performed preprocessing, dimensionality reduction and unsupervised clustering. Specifically, we used Seurat to log-normalize gene expression values, identify and select the top 2000 most highly variable genes, perform PCA linear dimensionality reduction, construct a KNN graph, and graph community-based unsupervised clustering. Next, we manually identified clusters corresponding to the isocortex. We then computed the gene expression signature score of our consensus gene signature using the ‘AddModuleScore’ function in Seurat, and performed linear regression of the signature score as a function of age in months separately for the wild-type and transgenic AD mouse models.

### Multiple hypothesis testing

In all cases, false discovery rate correction for multiple hypothesis testing was performed using the Benjamini-Hochberg procedure for FDR adjustment.

## Supporting information

Supplementary Table 4: Oligonucleotide sequences

## Data availability

All raw data have been deposited to NCBI’s GEO under accession GSE324610. All processed data have been deposited in the Single Cell Portal under accession SCP3562 (https://singlecell.broadinstitute.org/single_cell/study/SCP3562).

## Code availability

All code used for preprocessing SPACE-Tag data and reproducing the results in this manuscript have been deposited on GitHub at https://github.com/nygctech/SPACE-Tag/.

## Author contributions

C.Y. and S.V. conceived and designed the study with input from I.R. and discussions with S.M.; C.Y. and S.F. performed the experiments with help from M.P. and supervision from S.V.; R.H. and C.Y. analyzed the data with supervision from S.V.; S.F. annotated the histological sections; C.Y., R.H., S.F., M.P. and S.V. interpreted the data and wrote the manuscript with input from all the authors. All authors discussed the results.

## Acknowledgements

Work was supported by the Knut and Alice Wallenberg Foundation, Beijer Laboratory for Gene and Neuro Research, Swedish Society for Medical Research, Science for Life Laboratory, the MacMillan Center for the Non-Coding Genome, and 1RM1 HG011014-01 (S.V.). S.V. was supported as a Wallenberg Academy Fellow and SciLifeLab Fellow at Uppsala University.

## References

1. Sun, E. D. et al. Spatial transcriptomic clocks reveal cell proximity effects in brain ageing. Nature 638, 160–171 (2025).

2. Kang, S. H. et al. Degeneration and impaired regeneration of gray matter oligodendrocytes in amyotrophic lateral sclerosis. Nat. Neurosci. 16, 571–579 (2013).

3. Rozenblatt-Rosen, O. et al. Building a high-quality Human Cell Atlas. Nat. Biotechnol. 39, 149–153 (2021).

4. Oliver, A. J. et al. Single-cell integration reveals metaplasia in inflammatory gut diseases. Nature 635, 699–707 (2024).

5. Steyn, C. et al. A temporal cortex cell atlas highlights gene expression dynamics during human brain maturation. Nat. Genet. 56, 2718–2730 (2024).

6. Ounadjela, J. R. et al. Spatial multiomic landscape of the human placenta at molecular resolution. Nat. Med. 30, 3495–3508 (2024).

7. To, K. et al. A multi-omic atlas of human embryonic skeletal development. Nature 635, 657–667 (2024).

8. Zhang, D. et al. Spatial dynamics of brain development and neuroinflammation. Nature 647, 213–227 (2025).

9. Li, H. et al. Spatially decoding genotype-associated epigenetic landscapes in human lymphoma FFPE tissues via epi-Patho-DBiT. Nat. Commun. 17, 5924 (2026).

10. Deng, Y. et al. Spatial-CUT&Tag: Spatially resolved chromatin modification profiling at the cellular level. Science 375, 681–686 (2022).

11. Zhang, D. et al. Spatial epigenome–transcriptome co-profiling of mammalian tissues. Nature 1–10 (2023).

12. Guo, P. et al. Spatial profiling of chromatin accessibility in formalin-fixed paraffin-embedded tissues. Nat. Commun. 16, 5945 (2025).

13. Huang, Y.-H. et al. Unified molecular approach for spatial epigenome, transcriptome, and cell lineages. Proc. Natl. Acad. Sci. U. S. A. 122, e2424070122 (2025).

14. Llorens-Bobadilla, E. et al. Solid-phase capture and profiling of open chromatin by spatial ATAC. Nat. Biotechnol. 1–4 (2023).

15. Ståhl, P. L. et al. Visualization and analysis of gene expression in tissue sections by spatial transcriptomics. Science 353, 78–82 (2016).

16. Vickovic, S. et al. High-definition spatial transcriptomics for in situ tissue profiling. Nat. Methods 16, 987–990 (2019).

17. Kaya-Okur, H. S. et al. CUT&Tag for efficient epigenomic profiling of small samples and single cells. Nat. Commun. 10, 1930 (2019).

18. Fujii, T. et al. Reconstructing epigenomic dynamics through a single-cell multi-epigenome data integration framework. Nat. Commun. 16, 11006 (2025).

19. Bartlett, D. A. et al. High-throughput single-cell epigenomic profiling by targeted insertion of promoters (TIP-seq). J. Cell Biol. 220, (2021).

20. Harada, A. et al. A chromatin integration labelling method enables epigenomic profiling with lower input. Nat. Cell Biol. 21, 287–296 (2019).

21. Lu, T., Ang, C. E. & Zhuang, X. Spatially resolved epigenomic profiling of single cells in complex tissues. Cell 185, 4448–4464.e17 (2022).

22. Stickels, R. R. et al. Highly sensitive spatial transcriptomics at near-cellular resolution with Slide-seqV2. Nat. Biotechnol. 39, 313–319 (2021).

23. Liu, Y. et al. High-Spatial-Resolution Multi-Omics Sequencing via Deterministic Barcoding in Tissue. Cell 183, 1665–1681.e18 (2020).

24. Chen, A. et al. Spatiotemporal transcriptomic atlas of mouse organogenesis using DNA nanoball-patterned arrays. Cell https://doi.org/10.1016/j.cell.2022.04.003 (2022) doi:10.1016/j.cell.2022.04.003.

25. Chen, C. et al. Single-cell whole-genome analyses by Linear Amplification via Transposon Insertion (LIANTI). Science 356, 189–194 (2017).

26. Cusanovich, D. A. et al. A single-cell atlas of in vivo mammalian chromatin accessibility. Cell 174, 1309–1324.e18 (2018).

27. Zhong, C., Ang, K. S. & Chen, J. Interpretable spatially aware dimension reduction of spatial transcriptomics with STAMP. Nat. Methods 21, 2072–2083 (2024).

28. Calo, E. & Wysocka, J. Modification of enhancer chromatin: what, how, and why? Mol. Cell 49, 825–837 (2013).

29. Zhang, M. et al. Molecularly defined and spatially resolved cell atlas of the whole mouse brain. Nature 624, 343–354 (2023).

30. Alcamo, E. A. et al. Satb2 regulates callosal projection neuron identity in the developing cerebral cortex. Neuron 57, 364–377 (2008).

31. Lipiec, M. A. et al. TCF7L2 regulates postmitotic differentiation programmes and excitability patterns in the thalamus. Development 147, dev190181 (2020).

32. Bárcenas-Walls, J. R. et al. Nano-CUT&Tag for multimodal chromatin profiling at single-cell resolution. Nat. Protoc. 19, 791–830 (2024).

33. Zeisel, A. et al. Molecular architecture of the mouse nervous system. Cell 174, 999–1014.e22 (2018).

34. Okur, Z. et al. Control of neuronal excitation-inhibition balance by BMP-SMAD1 signalling. Nature 629, 402–409 (2024).

35. Apara, A. et al. KLF9 and JNK3 interact to suppress axon regeneration in the adult CNS. J. Neurosci. 37, 9632–9644 (2017).

36. Jobe, E. M. et al. Methyl-CpG-binding protein MBD1 regulates neuronal lineage commitment through maintaining adult neural stem cell identity. J. Neurosci. 37, 523–536 (2017).

37. Ho, W. Y. et al. TDP-43 mediates SREBF2-regulated gene expression required for oligodendrocyte myelination. J. Cell Biol. 220, e201910213 (2021).

38. Sen, P., Shah, P. P., Nativio, R. & Berger, S. L. Epigenetic Mechanisms of Longevity and Aging. Cell 166, 822–839 (2016).

39. Hahn, O. et al. Atlas of the aging mouse brain reveals white matter as vulnerable foci. Cell 186, 4117–4133.e22 (2023).

40. Raz, N., Ghisletta, P., Rodrigue, K. M., Kennedy, K. M. & Lindenberger, U. Trajectories of brain aging in middle-aged and older adults: regional and individual differences. Neuroimage 51, 501–511 (2010).

41. López-Otín, C., Blasco, M. A., Partridge, L., Serrano, M. & Kroemer, G. The hallmarks of aging. Cell 153, 1194–1217 (2013).

42. Bernstein, B. E., Meissner, A. & Lander, E. S. The mammalian epigenome. Cell 128, 669–681 (2007).

43. Yang, J.-H. et al. Loss of epigenetic information as a cause of mammalian aging. Cell 186, 305–326.e27 (2023).

44. Tong, H. et al. Quantifying the stochastic component of epigenetic aging. Nat. Aging 4, 886–901 (2024).

45. Meyer, D. H. & Schumacher, B. Aging clocks based on accumulating stochastic variation. Nat. Aging 4, 871–885 (2024).

46. Ernst, J. & Kellis, M. ChromHMM: automating chromatin-state discovery and characterization. Nat. Methods 9, 215–216 (2012).

47. Wang, L. et al. Spatial transcriptomics of the aging mouse brain reveals origins of inflammation in the white matter. Nat. Commun. 16, 3231 (2025).

48. Gao, Y. et al. Continuous cell-type diversification in mouse visual cortex development. Nature 647, 127–142 (2025).

49. Multiomic Integration Neuroscience Application Note: Visium for FFPE Plus Immunofluorescence Alzheimer’s Disease Mouse Model Brain Coronal Sections from One Hemisphere Over a Time Course. 10x Genomics https://www.10xgenomics.com/datasets/multiomic-integration-neuroscience-application-note-visium-for-ffpe-plus-immunofluorescence-alzheimers-disease-mouse-model-brain-coronal-sections-from-one-hemisphere-over-a-time-course-1-standard.

50. Silverman, E. S. et al. cAMP-response-element-binding-protein-binding protein (CBP) and p300 are transcriptional co-activators of early growth response factor-1 (Egr-1). Biochem. J. 336 ( **Pt 1**), 183–189 (1998).

51. Lai, B. et al. MLL3/MLL4 are required for CBP/p300 binding on enhancers and super-enhancer formation in brown adipogenesis. Nucleic Acids Res. 45, 6388–6403 (2017).

52. Brooks, Y. S. et al. Multifactorial ERβ and NOTCH1 control of squamous differentiation and cancer. J. Clin. Invest. 124, 2260–2276 (2014).

53. Moses, L. & Pachter, L. Museum of spatial transcriptomics. Nat. Methods 19, 534–546 (2022).

54. Vickovic, S. et al. Massive and parallel expression profiling using microarrayed single-cell sequencing. Nat Commun 7, 13182 (2016).

55. Martin, M. Cutadapt removes adapter sequences from high-throughput sequencing reads. EMBnet J. 17, 10 (2011).

56. Smith, T., Heger, A. & Sudbery, I. UMI-tools: modeling sequencing errors in Unique Molecular Identifiers to improve quantification accuracy. Genome Res. 27, 491–499 (2017).

57. Langmead, B. & Salzberg, S. L. Fast gapped-read alignment with Bowtie 2. Nat. Methods 9, 357–359 (2012).

58. Zhang, Y. et al. Model-based analysis of ChIP-Seq (MACS). Genome Biol. 9, R137 (2008).

59. Daley, T. & Smith, A. D. Predicting the molecular complexity of sequencing libraries. Nat. Methods 10, 325–327 (2013).

60. Stuart, T., Srivastava, A., Madad, S., Lareau, C. A. & Satija, R. Single-cell chromatin state analysis with Signac. Nat. Methods 18, 1333–1341 (2021).

61. Maniatis, S. et al. Spatiotemporal dynamics of molecular pathology in amyotrophic lateral sclerosis. Science 364, 89–93 (2019).

62. Ramírez, F. et al. deepTools2: a next generation web server for deep-sequencing data analysis. Nucleic Acids Res. 44, W160–5 (2016).

63. Guo, P. et al. Multiplexed spatial mapping of chromatin features, transcriptome and proteins in tissues. Nat. Methods 1–10 (2025).

64. DeTomaso, D. & Yosef, N. Hotspot identifies informative gene modules across modalities of single-cell genomics. Cell Syst. 12, 446–456.e9 (2021).

65. Whyte, W. A. et al. Master transcription factors and mediator establish super-enhancers at key cell identity genes. Cell 153, 307–319 (2013).

66. Wang, Y. et al. SEdb 2.0: a comprehensive super-enhancer database of human and mouse. Nucleic Acids Res. 51, D280–D290 (2023).

67. Weirauch, M. T. et al. Determination and inference of eukaryotic transcription factor sequence specificity. Cell 158, 1431–1443 (2014).

68. Ekvall, M. et al. Spatial landmark detection and tissue registration with deep learning. Nat. Methods 21, 673–679 (2024).

69. Heumos, L. et al. Best practices for single-cell analysis across modalities. Nature Reviews Genetics 24, 550–572 (2023).

70. Kraskov, Alexander and St\"ogbauer, Harald and Grassberger, Peter. Estimating mutual information. Phys. Rev. E 69, 066138 (2004).

71. Ma, J. Copent: Estimating copula entropy and transfer entropy in R. arXiv [stat.CO] (2020).

72. Amemiya, H. M., Kundaje, A. & Boyle, A. P. The ENCODE Blacklist: Identification of Problematic Regions of the Genome. Scientific Reports 9, 9354 (2019).

73. Peidli, S. et al. scPerturb: harmonized single-cell perturbation data. Nature Methods 21, 531–540 (2024).

74. Bartosovic, M., Kabbe, M. & Castelo-Branco, G. Single-cell CUT&Tag profiles histone modifications and transcription factors in complex tissues. Nat. Biotechnol. 39, 825–835 (2021).

