## Supplemental Data 1 for "SPACE-Tag Enables High-Throughput Spatial Chromatin Profiling With CUT&Tag"

Contents: Supplementary Figures 1–14. Supplementary Table 4 (oligonucleotide sequences) is provided as a separate file; Supplementary Tables 1–3 will be provided with the published version of this article. Reference numbers in the legends refer to the main-text reference list.

### Supplementary Figures

**Supplementary figure 1**

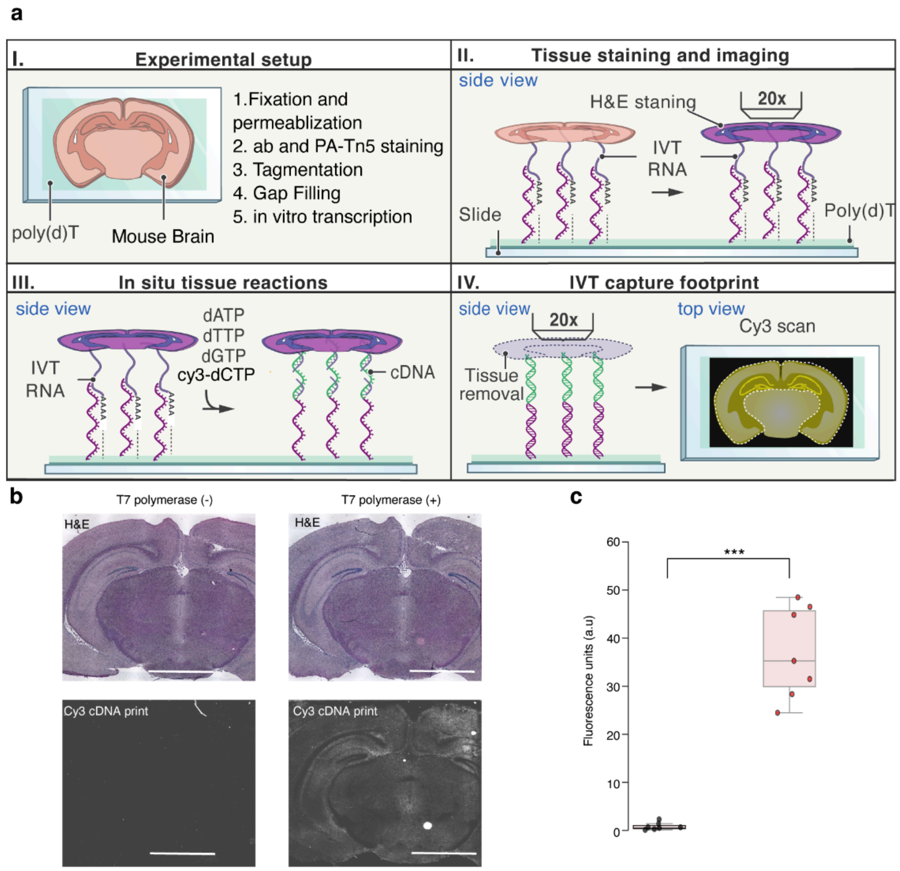


**Supplementary Figure 1**. **Effect of T7 amplification on signal capture in SPACE-Tag.** (**a**) Schematic of the workflow for signal quantification. (I) Brain sections are deposited on a pseudospatial array with a poly(dT) capture area, stained for primary and secondary antibodies, and *in situ* tissue reactions are performed: pA-Tn5 tethering, tagmentation, gap-filling and HCl permeabilization. The SPACE-Tag procedure then utilizes T7-based amplification to generate amplified RNA molecules that are subsequently captured on the pseudospatial slide. (II) HE histology staining was performed and imaged. (III) Fluorescent cDNA molecules are then generated with a reverse transcription reaction using fluorescently labeled nucleotides. (IV) Tissue was removed from the surface and fluorescent signal was captured through fluorescence imaging. (**b**) H&E images (top) and cDNA signal intensity (bottom) of sections with (T7 (+)) or without (T7 (-)) the T7 amplification step. Scale bars are 400 µm. Bright pixels correspond to higher intensity signals. (**c**) Quantification of the fluorescence signals from (**b**). For each anatomical region (hippocampus, isocortex, and hypothalamus), signal intensities from three regions of interest (ROIs) were analyzed across seven independent tissue sections per condition (T7 polymerase (-) and T7 polymerase (+)). Dots: Individual samples (n). Error bars represent s.e.m. Significance testing: Wilcoxon's rank sum test; *** denotes p<0.001.

**Supplementary figure 2**

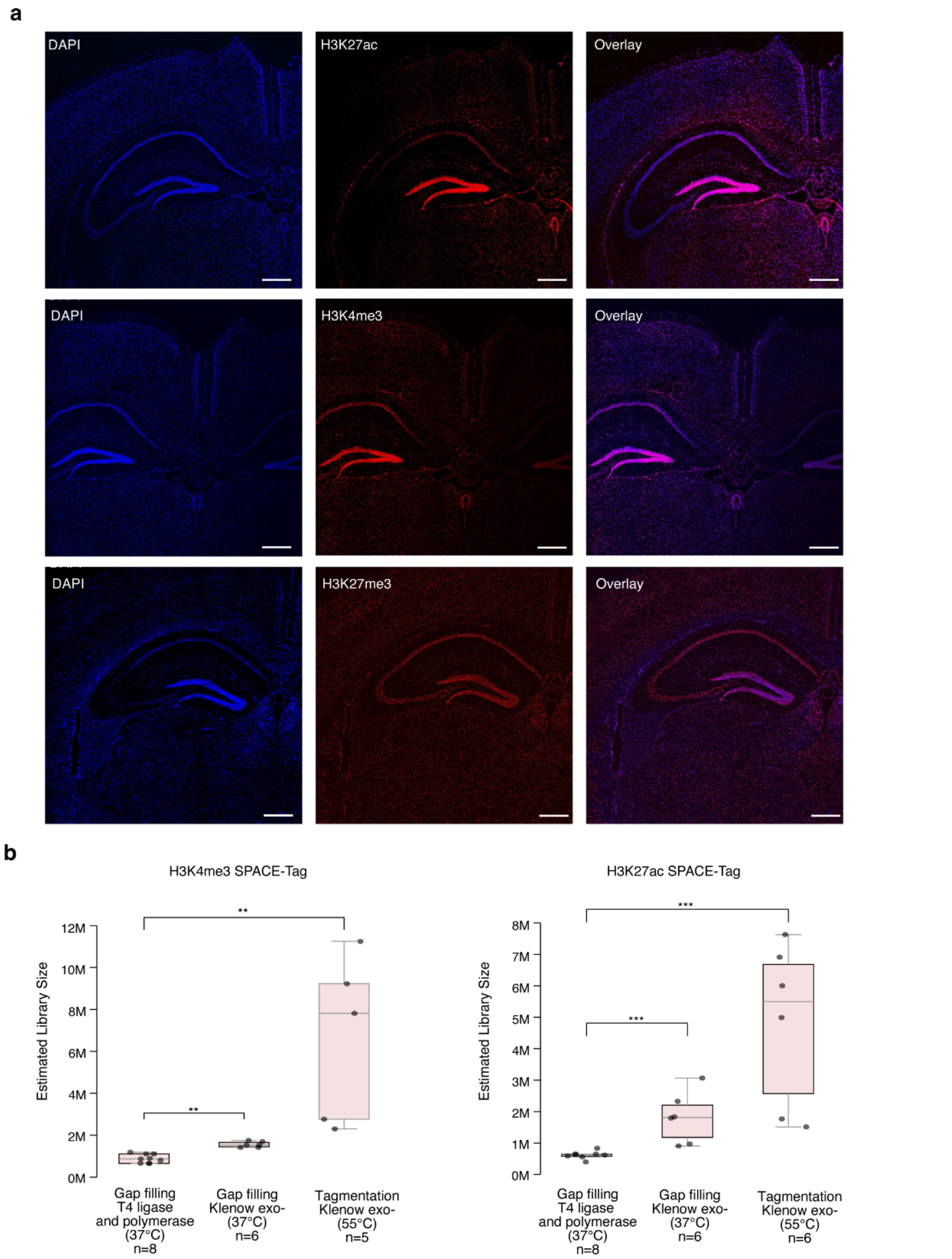


**Supplementary Figure 2. Optimized antibody staining, gap-filling and tagmentation in adult mouse brain sections.** (**a**) Fluorescence images of nuclear (DAPI; blue) and antibody stains (red) for H3K27ac (top), H3K4me3 (middle) and H3K27me3 (bottom). Channels overlay shown on the right. Scale bar: 400 µm. (**b**) Library complexity across protocol optimizations. Estimated number of unique fragments (Methods) in the resulting cDNA libraries (y-axis) as a function of experimental parameters. Comparisons include gap-filling and ligation chemistry (T4 DNA Polymerase/Ligase vs. Klenow Polymerase) and tagmentation temperature (37 °C vs. 55 °C). Data are shown for the active marks H3K4me3 (left) and H3K27ac (right). Gap-filling using Klenow polymerase (mid) significantly increased the library complexity compared to T4 polymerase and ligase (left).. Additionally, performing the tagmentation reaction at 55°C (right) further enhanced complexity relative to the conventional 37°C reaction temperature (mid). Dots: Individual samples (n). Error bars represent s.e.m. Significance testing: Wilcoxon’s rank sum test; ** denotes $p<0.01$; *** denotes $p<0.001$.

Supplementary figure 3

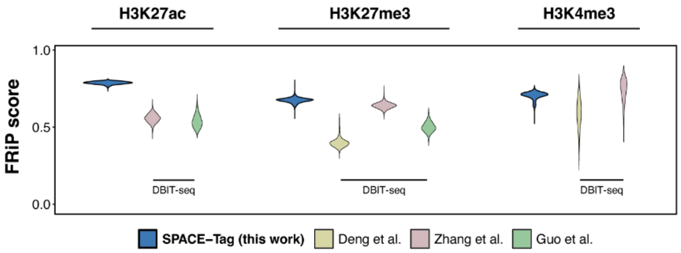


**Supplementary Figure 3. Comparing SPACE-Tag against existing spatial CUT&Tag technologies.** Distributions of Fraction of Reads in Peaks (FRiP) values (y-axis) across SPACE-Tag and indicated baseline methods (color code). Higher scores indicate superior enrichment of histone marks within identified peaks.

**Supplementary figure 4**

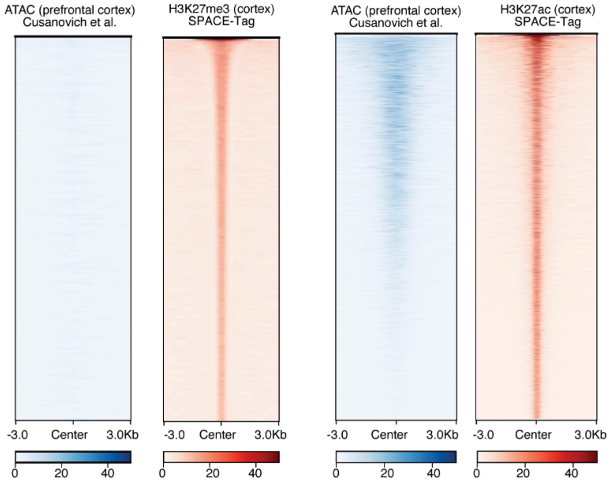


**Supplementary Figure 4. Specificity of SPACE-Tag signal relative to chromatin accessibility.** Comparison of ATAC-seq signal intensity at repressive (H3K27me3, left) versus active (H3K27ac, right) peaks defined by SPACE-Tag. The lack of ATAC-seq signal at repressive marks underscores the specificity of epitope capture, while the strong overlap with H3K27ac validates the capture of functionally active regulatory elements.

**Supplementary figure 5**

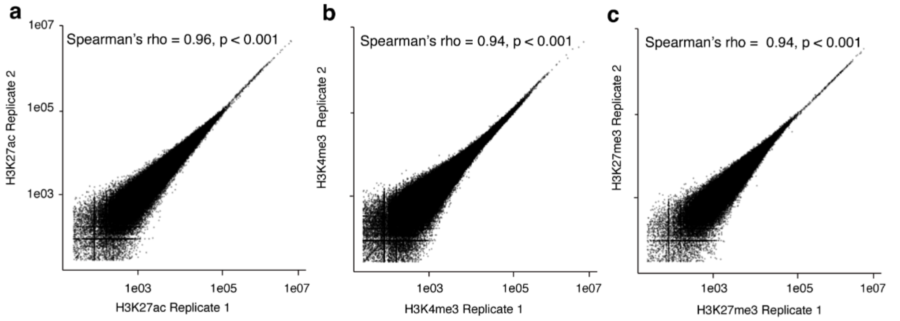


**Supplementary Figure 5. SPACE-Tag reproducibility.** (**a-c**) Reproducibility of SPACE-Tag experiments across biological replicates for (**a**) H3K27ac, (**b**) H3K4me3 and (**c**) H3K27me3. The genome was split into 10 kb tiles, and a scatterplot of normalized SPACE-Tag signals between replicate experiments was plotted for all tiles (**Methods**). The Spearman correlation coefficient was calculated for each modification; significance was determined by a two-sided *t*-test ($p<0.001$ denoted).

**Supplementary figure 6**

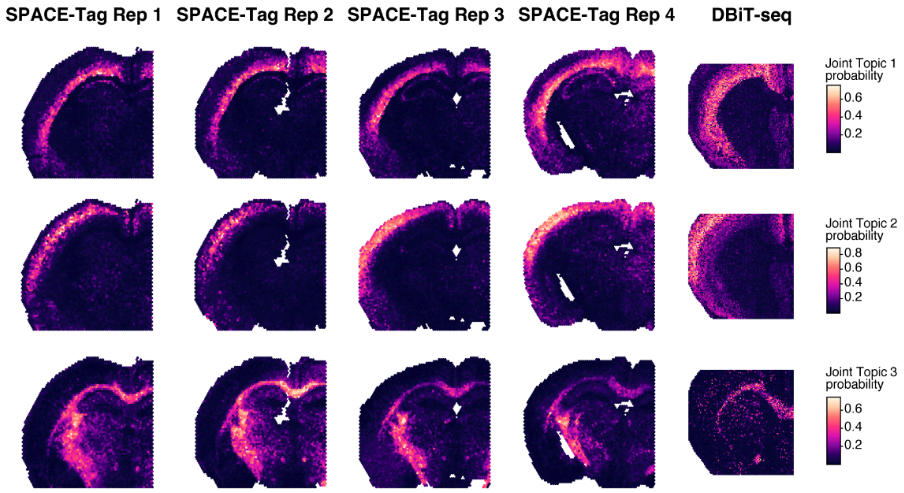


**Supplementary Figure 6. Sensitivity comparison in discovering spatial domain between SPACE-Tag and DBiT-seq.** Joint topic modelling of 4 replicate SPACE-Tag sections and 1 DBiT-seq spatial CUT&Tag section of H3K27ac profiling in mouse brain. Each row represents a single inferred joint topic; color scale represents topic probability.

Supplementary figure 7

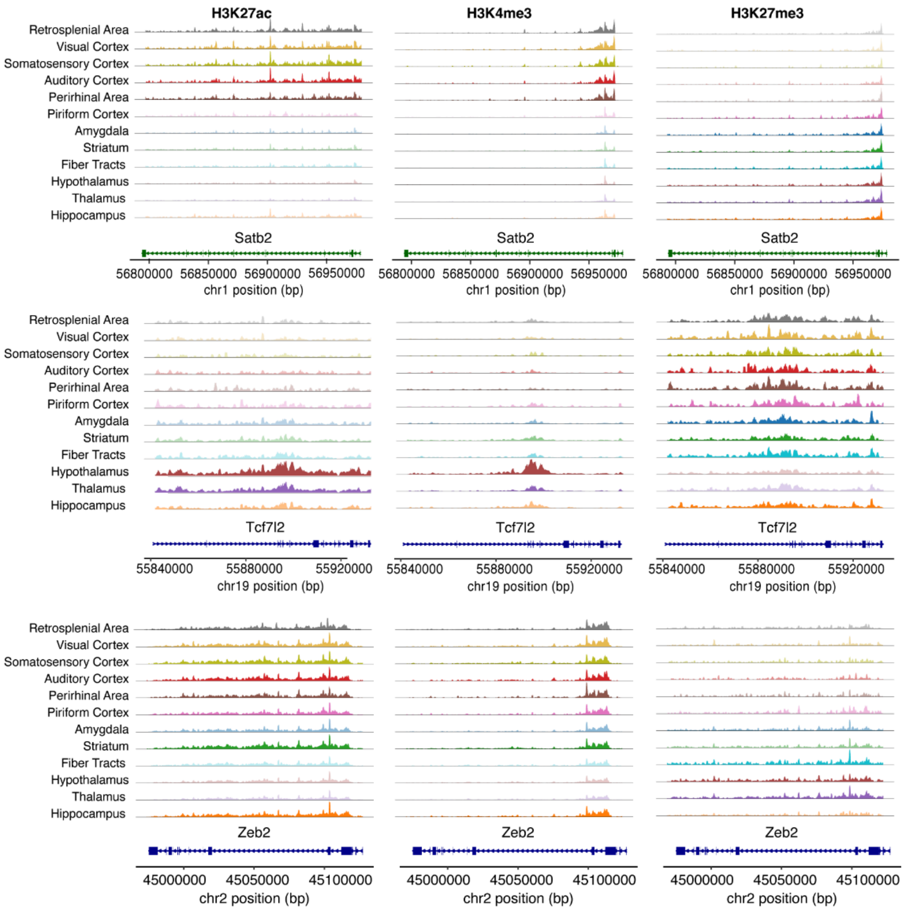


**Supplementary Figure 7. Genome coverage tracks of region-specific key neuronal marker genes profiled by SPACE-Tag.** Genome coverage tracks showing normalized epigenomic signal (y-axis; left: H3K27ac; middle: H3K4me3; right: H3K27me3) across the gene body and flanking regions (x-axis) of marker genes (top: *Satb2*, middle: *Tcf7l2*; bottom: *Zeb2*). The corresponding gene body is depicted beneath each track, along with genomic coordinates. Tracks for tissue regions with high expected signal in each mark are highlighted in bold.

Supplementary figure 8

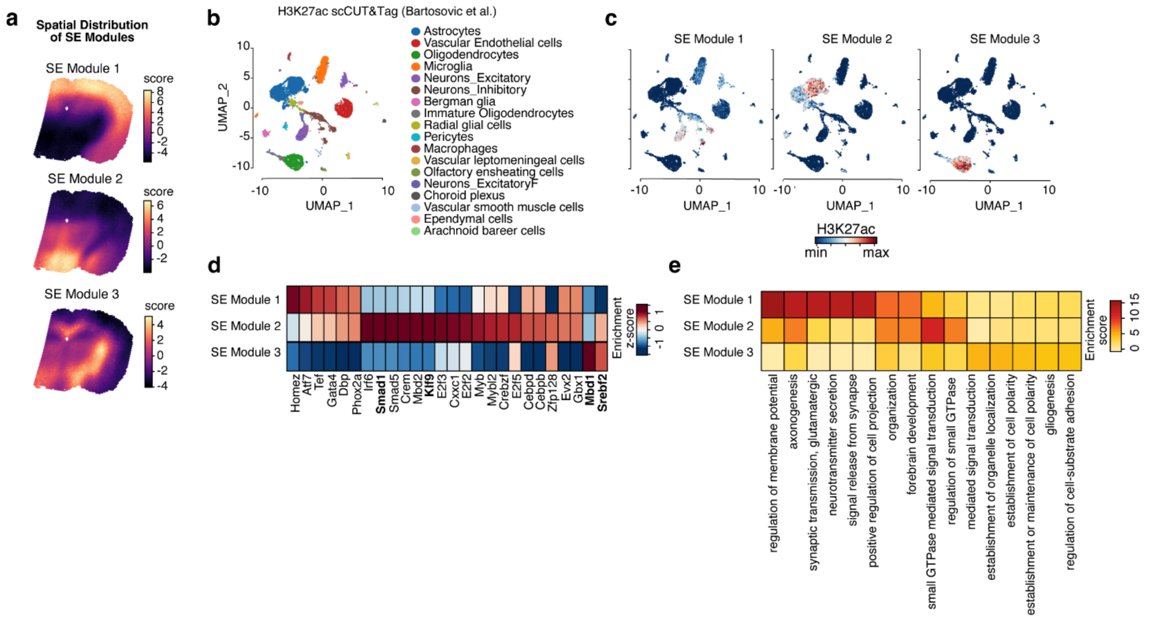


**Supplementary Figure 8. Super-enhancer modules are active in specific regions and cell types.** (**a**) Spatial distribution (color scale) of identified SE modules across brain sections. Module 1 occupies the cortex regions. Module 2 occupies the hypothalamus region. Module 3 occupies white matter tracts. **(b)** UMAP embedding of cell types derived from a previously published H3K27ac single-cell CUT&Tag dataset of the postnatal mouse brain[74]. (**c**) H3K27ac signal (color scale) of SE modules (columns) across cell types (color scale); Module 1 (left) is primarily enriched in excitatory and inhibitory neurons, Module 2 (middle) in astrocytes, and Module 3 (right) in oligodendrocytes. (**d, e**) Enrichment scores (color scale) of (**d**) differentially enriched TF motifs (columns) and (**e**) GO terms within each SE module (rows).

Supplementary figure 9

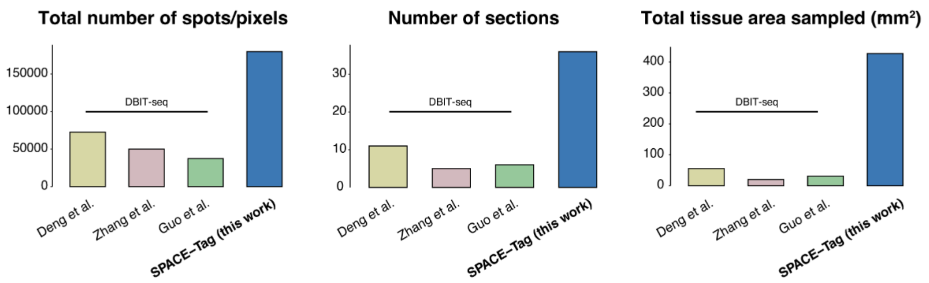


**Supplementary Figure 9. SPACE-Tag sample numbers.** Comparison of the SPACE-Tag dataset generated in this manuscript with existing spatial CUT&Tag studies highlights the scalability of SPACE-Tag. Left: total number of spots (SPACE-Tag) or pixels (DBiT-seq) profiled per study. Middle: unique number of sections profiled per study. Right: total tissue area profiled under spots/pixels per study.

**Supplementary figure 10**

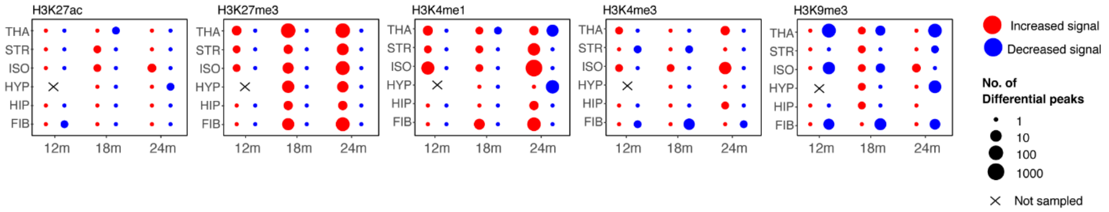


**Supplementary Figure 10. Regional shifts in histone mark occupancy during aging.** Comparison of histone peak signals across various brain regions at 12m, 18m and 24m relative to 3m. Dot size represents the number of significant differentially enriched peaks, and color indicates the direction of change. Results highlight the region-specific nature of age-associated epigenetic remodeling.

Supplementary figure 11

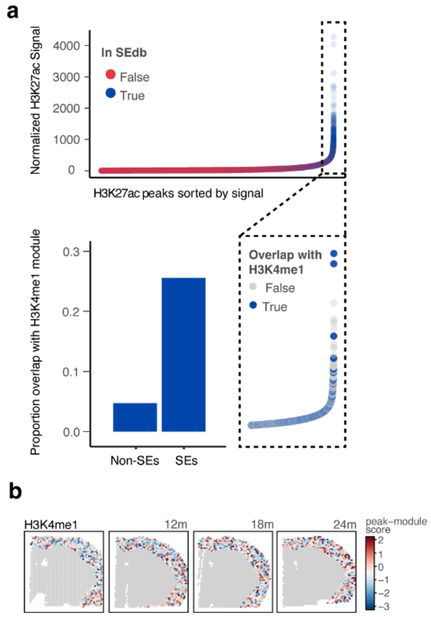


**Supplementary Figure 11. Preferential enrichment of the age-associated H3K4me1 module at isocortex super-enhancers. (a)** Top panel: dotplot of normalized signal intensity at isocortex-specific H3K27ac peaks, categorized by overlap with the SEdb database (blue) or not (red). **Bottom right:** Magnified view of high-signal H3K27ac peaks classified as SEs. Blue color indicates peaks overlapping with the isocortex-specific H3K4me1 peak module, highlighting the expansion of this mark at these regulatory hubs. **Bottom left:** Quantitative analysis showing the fraction of SEs versus non-SE enhancers overlapping with the isocortex-specific H3K4me1 peak module. These data reveal a significant bias for age-dependent H3K4me1 expansion at SE regions. (**b**) Spatial plots of H3K27ac scores at isocortex peak module across mouse brain aging. Color represents normalized module signal.

**Supplementary figure 12**

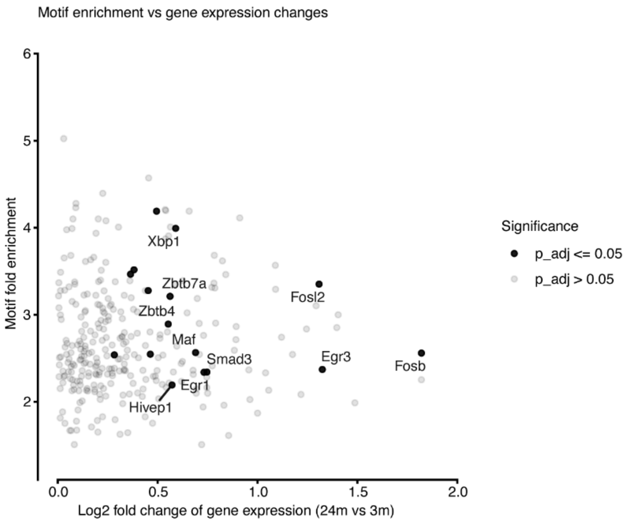


**Supplementary Figure 12. Identification of TFs exhibiting significant motif enrichment and differential gene expression.** Scatterplot of motif fold enrichment (y-axis) versus differential gene expression (x-axis) across mouse TFs. Solid points represent TFs with significant differential RNA expression.

**Supplementary figure 13**

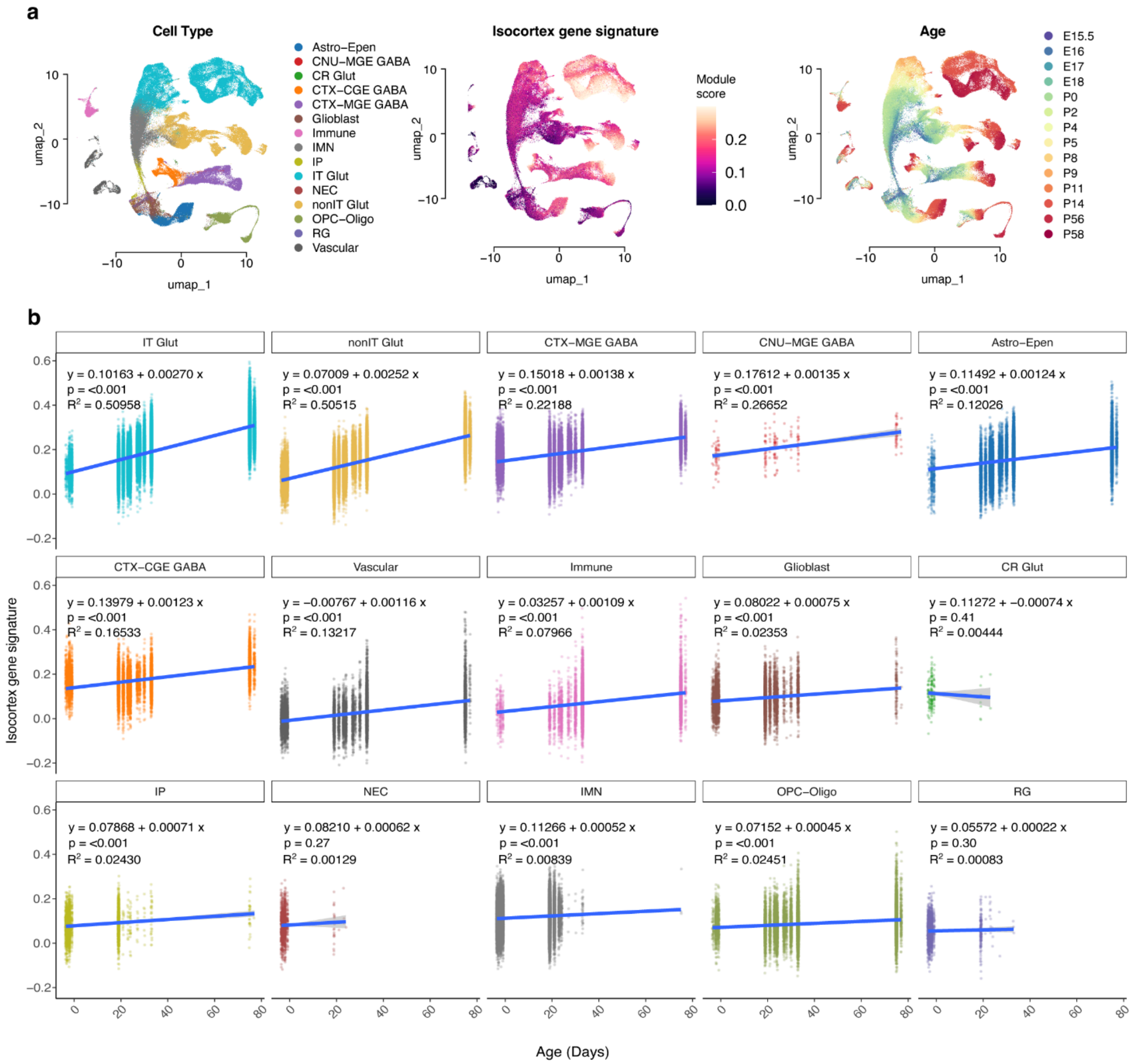


**Supplementary Figure 13. Cross-referencing the consensus isocortex gene signature during mouse brain development.** (**a**) Left: UMAP of single-nucleus RNA-seq[48], colored by cell type annotation. Middle: Consensus isocortex gene signature during mouse development. Color represents normalized signature expression. Right: UMAP of snRNA-seq data colored by developmental stage. (**b**) Cell-type-specific expression of the isocortex consensus gene signature across time. Y-axis represents normalized signature expression, X-axis represents age in days. Solid blue line represents fitted linear regression line.

**Supplementary figure 14**

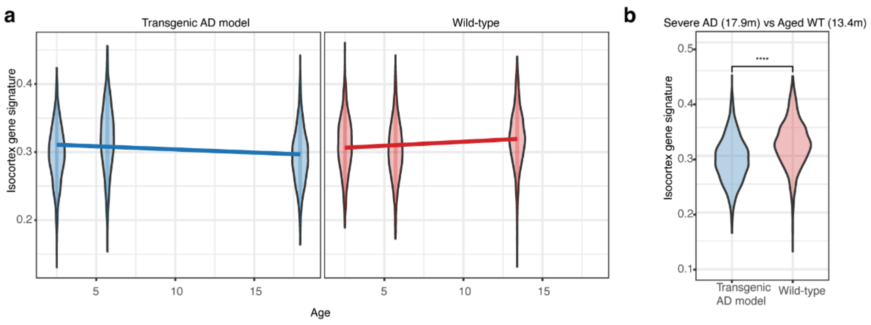


**Supplementary Figure 14. Cross-referencing the consensus isocortex gene signature in the AD mouse model.** (**a**) Expression of isocortex consensus gene signature in isocortex Visium transcriptomics data of (right) wild-type and (left) transgenic AD mouse models over time. Y-axis represents normalized signature expression, X-axis represents age in months. Solid line represents fitted linear regression line. (**b**) Violin plot comparing consensus signature expression between aged wild-type and severe AD conditions. Annotation represents significance by one-sided Wilcoxon rank-sum test; **** denotes $p < 0.0001$.

**Supplementary Tables**
**Supplementary Table 1.** SEs associated with spatial distributions and cell types.
**Supplementary Table 2.** DE genes associated with region-specific aging.
**Supplementary Table 3.** Isocortex genes associated with chromatin changes.
**Supplementary Table 4.** Oligonucleotide primer sequences.
